# Mammalian enhancers and GWASs act proximally and seldom skip active genes

**DOI:** 10.1101/2024.11.29.625864

**Authors:** Takeo Narita, Chunaram Choudhary

## Abstract

Enhancers play a critical role in regulating transcription. Nearly 90% of human genetic variants identified in genome-wide association studies (GWAS) are located in distal regions, underscoring the importance of enhancers in human development, diseases, and traits. It is widely suggested that mammalian enhancers frequently skip active genes, and thus, linear proximity is a poor predictor of their targets. A key unresolved question is how often mammalian enhancers skip proximal active genes to specifically target distal genes. Genome-wide enhancer-promoter mapping shows that enhancers frequently bypass active genes, while ultra-deep locus-specific analyses reveal extensive multi-way interactions between enhancers and promoters, forming nested microcompartments. The functional significance of these seemingly contrasting phenomena remains unclear.

Here, we compared hundreds of enhancer-target gene pairs identified using enhancer-promoter chromatin contact maps, enhancer-promoter RNA interaction data, and genome-scale CRISPR interference (CRISPRi) perturbations. Our findings reveal limited overlap between active gene-skipping enhancer-gene pairs identified through physical interaction mapping and CRISPRi. Additionally, promoters involved in multi-way enhancer interactions are not co-regulated by shared coactivators. Notably, gene-skipping and non-skipping enhancers identified via CRISPRi differ fundamentally in chromatin features, gene activation strength, false discovery rates, target gene distance, coactivator requirements, and cell-type specificity of target genes. These results suggest that gene-skipping enhancer-promoter interactions observed in chromatin and RNA-based analyses do not reliably predict functional enhancer-gene relationships. We propose that linear enhancer-promoter proximity and coactivator dependency offer a simple, scalable, and cost-effective method for genome-wide prediction of enhancer and GWAS targets, with accuracy comparable to state-of-the-art experimental techniques. While enhancers can skip active genes, such deliberate skipping appears to be the exception rather than the rule.

## Introduction

Enhancers are crucial transcription regulators in metazoans ^1–3^, housing nearly 90% of human genetic variants identified in genome-wide association studies (GWAS) ^4–6^. They function in an orientation-independent manner and can act over long distances to activate distal targets ^1–3,7^. Mammalian genomes are organized into loops and topologically-associating domains, facilitating interactions between distant cis-regulatory elements ^8–10^. Because of this intricate 3D genome organization, enhancers can skip proximal genes to activate distant ones, and multiple enhancers can activate one or more genes. For example, enhancers regulating *Shh*, *Igf2*, and alpha-globin genes skip over an active gene ^11–13^, while multiple enhancers converge to activate multiple genes in the beta-globin gene cluster ^14^.

Their ability to act at long distances and skip genes poses a challenge in identifying enhancer and GWAS target genes. To address this, various chromosome conformation capture (3C)-based methods, such as HiC, Micro-C, Hi-ChIP, PC-HiC, ChIA-PET, SPRITE, and GAM, have been used to map enhancer-promoter (E-P) contacts (or E-P loops) ^10,15–27^. Genome-scale E-P contact maps generated by these methods show that a large portion, often the majority, of enhancers skip nearby genes to target distal ones ^10,15–27^. For instance, an early analysis by ENCODE found that only ∼7% of chromatin looping interactions are with the nearest gene ^18^. Since then, many studies have confirmed this observation using different 3C-based techniques ^10,15–27^. Recent RNA in situ conformation sequencing (RIC-seq)-based E-P mapping also shows that 87% of human enhancers avoid the nearest genes and interact with distal promoters ^28^. Indeed, going beyond intra-chromosomal gene skipping, a large portion of enhancers interact with promoters on other chromosomes ^15,28–32^.

As the depth and resolution of genome contact mapping increase, the complexity of E-P interactions is becoming ever more complex. Ultra-deep, locus-specific analyses reveal that enhancers and promoters engage in multi-way interactions and form nested microcompartments ^33–35^, and DNA ligation-free methods suggest that the prevalence of multi-way interactions in active chromatin regions is underestimated in traditional approaches like Hi-C^36^.

Based on the covariance in E-P interactions and gene expression, an increasing number of efforts directly link chromatin contacts with gene regulation. Within the past year, several single-cell-based methods, such as HiRES ^31^, MUSIC ^37^, LiMCA ^38^, and GAGE-seq ^39^, have been developed for this. A major conclusion of these single-cell-based analyses is that a large fraction of enhancers deliberately skips active genes to activate distal targets. For example, single-cell HiRES analyses imply that nearly 90% of cell type-specific differential interactions affect distal targets, and a large portion exceeds >1Mb distance ^31^. A limitation of these approaches is that E-P relationships are inferred indirectly from the covariation in E-P contacts and gene expression. Functional analysis of enhancers by genome-wide single-cell CRISPR interference (CRISPRi) shows that one-third of human enhancers activate non-nearest genes within 1Mb of targeted enhancers ^40^.

Although E-P interactions are being mapped at an extraordinary resolution and scale, several key questions remain unanswered:

- Is linear or 3D proximity a better predictor of functional relationships?
- Do all physical E-P contacts inform about functionally relevant gene regulatory interactions? If not, what distinguishes functional from non-functional E-P contacts?
- In multi-way E-P interactions, are all interacting promoters similarly activated by enhancers?
- Do gene-skipping and non-skipping enhancers identified in global CRISPRi analyses share the same attributes, function through the same mechanisms, and exert similar effects on gene regulation?

We systematically analyzed the properties of E-P pairs identified by Micro-C, RIC-seq, and CRISPRi, revealing striking differences between gene-skipping and non-skipping enhancers. Contrary to prevailing models, our findings suggest that enhancers rarely skip active genes. Instead, they primarily act proximally and regulate genes through a unified mechanism.

## Results

### Datasets for comparing properties of E-P pairs

To identify general enhancer properties, we analyzed E-P pairs identified using three distinct methods by four research groups, all conducted in the same human lymphoblast cell line, K562.

(1) Enhancer-gene (E-G) pairs identified by genome-wide, single-cell CRISPRi ^40^ were used as the reference dataset for experimentally defined enhancers and their target genes.
(2) A separately generated, focused (not genome-scale) CRISPRi dataset ^41^ was used to confirm the properties of E-G pairs discovered in the genome-scale CRISPRi dataset.
(3) Micro-C mapped E-P contacts ^42^ served as a representative dataset for assessing the properties of E-P pairs identified by 3C-based methods.
(4) RIC-seq ^28^ identified E-P pairs were used to identify properties of enhancer targets identified by RNA-based interactions, distinct from chromatin-independent approaches.

Additionally, we analyzed co-regulation of promoters showing multi-way interactions in locus-specific analyses ^34^ and examined co-activator dependency of genes proximal to MED1-enriched super-enhancers ^43^ in mouse embryonic stem cells (mESC). For clarity, unless specified otherwise, throughout this work, the word ‘enhancer’ refers to candidate enhancers, and ‘enhancer target’ refers to candidate enhancer-regulated genes.

### Gene-skipping E-P pairs identified by Micro-C and CRISPRi overlap minimally

We used unbiased, genome-scale, single-cell CRISPRi screen data ^40^ as the reference dataset of experimentally defined functional E-P pairs. This dataset encompassed 664 functional E-G pairs, including 470 high-confidence E-G (^highconf^E-G) and 194 low-confidence E-G (^lowconf^E-G) pairs, involving 441 and 182 enhancers, respectively.

Our primary goal was to understand the properties of gene-skipping and non-skipping enhancers. Therefore, CRISPRi targeted enhancers were grouped into active gene ‘skipping’ and ‘non-skipping.’ In the analyses by Gasperini et al., enhancers not impacting the nearest active gene were considered as ‘skipping’; from this definition, approximately one-third of enhancers skip active genes^40^. We considered that an enhancer could activate one or both flanking genes without ‘skipping over’ an intervening active promoter. To account for this, we classified enhancers linked to one or both actively transcribed immediate neighbor genes as ‘non-skipping,’ while those linked to non-neighboring active genes as ‘skipping’ (See Methods). According to this criterion, 25.9% of E-G pairs involved skipping of active genes, and the proportion of gene-skipping E-G pairs was more than twice in ^lowconf^E-G pairs compared to ^highconf^E-G pairs (43.8% versus 18.5%) (**Supplemental Figure 1A**).

Among genome-scale chromatin contact mapping methods, Micro-C can map contacts at high resolution ^16,44^. Therefore, we used Micro-C data ^42^ for our analyses. Micro-C detected chromatin loops were called using HiCCUPS ^45^ at 1kb, 2kb, and 5kb bin resolutions, and the identified loops from the three resolutions were merged. If loops were detected at multiple resolutions, only those from the higher resolution were retained. Micro-C loops were considered to overlap with CRISPRi-identified E-G pairs if they matched within a 1kb distance. CRISPRi analyzed 78,776 E-G pairs involving 5,779 enhancers and their proximal genes within 1 Mb (**Figure 1A**), and from them, Micro-C detected loops between 885 E-P pairs, 57% non-skipping and 43% gene-skipping (**Figure 1B**).

**Figure 1:**
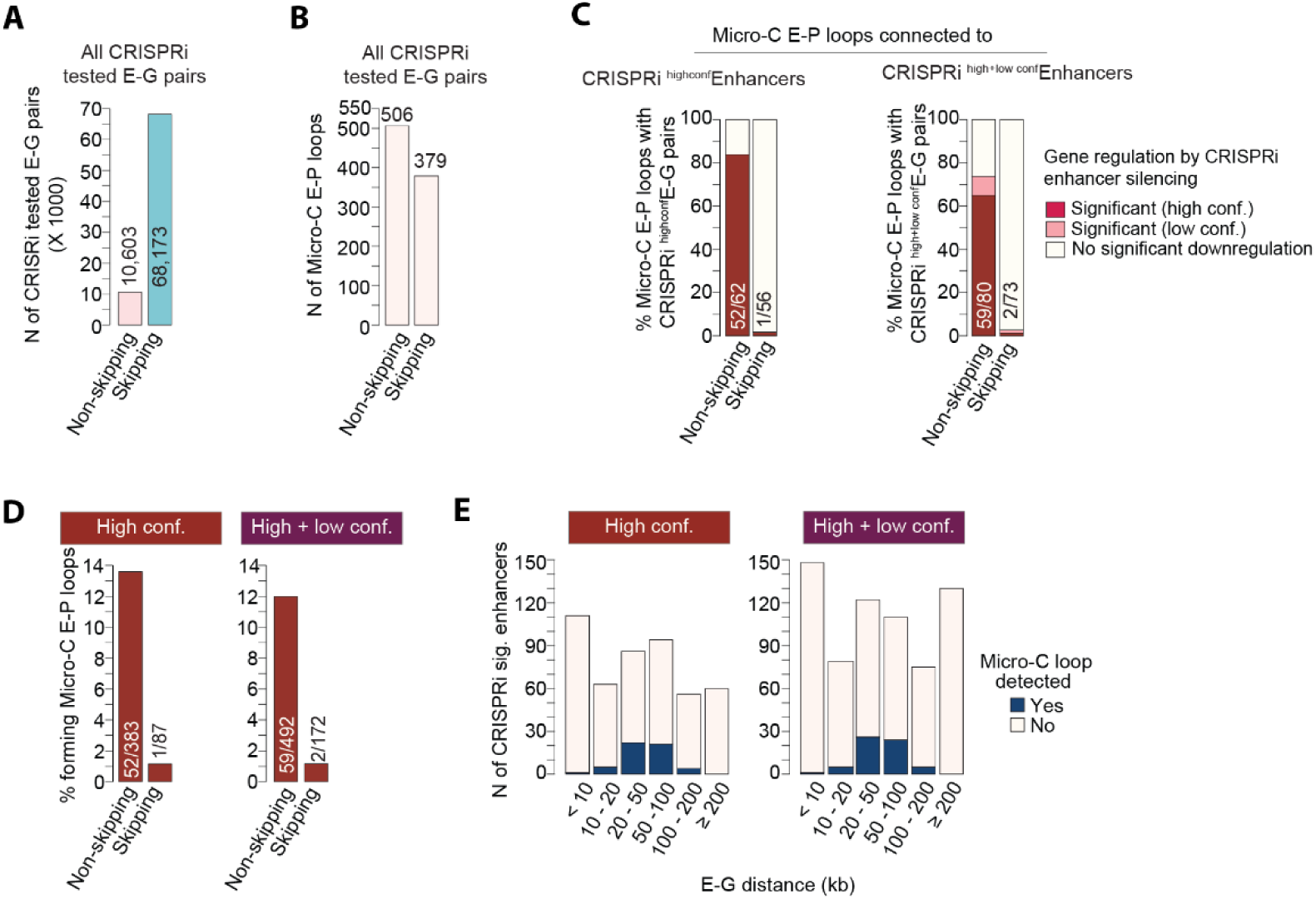
Distinct overlap between non-skipping and gene-skipping E-P pairs identified by Micro-C and CRISPRi. (**A-B**), The number of CRISPRi-tested E-G pairs within 1 Mb of the targeted enhancers (**A**), and the number of CRISPRi-tested E-G pairs showing chromatin interaction in Micro-C data (**B**). E-G pairs are classified as active gene-skipping and non-skipping (see Methods). (**C**) Micro-C identified E-P loops involving CRISPRi-defined significant enhancers. E-P loops involving high-confidence (high-conf.) and high + low confidence (high+low conf.) enhancers are analyzed separately. Micro-C-identified E-P pairs are grouped as active gene-skipping and non-skipping. Within each group, regulation of enhancer-interacting genes after CRISPRi enhancer silencing is indicated. (**D**) CRISPRi identified significant E-G pairs grouped into gene-skipping and non-skipping groups, and the percent of pairs showing E-P looping in Micro-C is shown. (**E**) CRISPRi identified significant E-G pairs grouped into indicated distance categories, and within each category, the number of pairs with or without detectable E-P looping in Micro-C is shown.

We focused on E-P loops that involve CRISPR-defined functional enhancers. If the corresponding genes in Micro-C identified E-P loops are identified as enhancer targets by CRISPRi, the functional relationship between the Micro-C identified E-P loops will be validated. If the genes in Micro-C identified E-P loops are not identified as enhancer targets by CRISPRi, it would imply that Micro-C E-P loops are inadequate to inform on functional E-P relationship or CRISPRi failed to identify those enhancer targets.

Micro-C identified 153 E-P loops, which involved CRISPRi-confirmed functional enhancers, and for which data are available for the regulation of the contacting genes after CRISPRi silencing the contacting enhancers (**Figure 1C**). Among these, the corresponding genes in the gene-skipping and non-skipping E-P loops show notable differences in the regulation by CRISPRi (**Figure 1C**). While most of the corresponding genes in non-skipping E-P loops are confirmed as genuine enhancer targets by CRISPRi, most genes in gene-skipping E-P loops are not.

Among CRISPRi-validated E-G pairs, Micro-C loops were almost exclusively detected for non-skipping pairs. Of 172 CRISPRi-identified gene-skipping functional E-G pairs, Micro-C confirmed looping between just two (**Figure 1D**). Micro-C mostly detected looping between E-P pairs with 20-100kb distances (**Figure 1E**). Among E-G pairs with 20-100kb distance, Micro-C identified looping between >22% of them. In contrast, few or not loops were identified for E-G pairs with <10KB or >200 kb distances.

This shows that CRISPRi confirms the functional relationship for most non-skipping E-P loops identified by Micro-C but seldom confirms it for gene-skipping E-P loops. Conversely, Micro-C detects looping between CRISPRi-identified non-skipping E-G pairs but rarely detects looping between E-G pairs that involve gene skipping.

### Histone H2B acetylation is predictive of functional enhancers

The results raise the question: Why is looping not detected between CRISPRi-identified E-G pairs involving active gene skipping? Is this solely due to the longer distances of gene-skipping E-G pairs and the technical limitations of Micro-C in identifying these long loops? Or could it be that CRISPRi-identified gene-skipping and non-skipping enhancers have distinct attributes?

To address these questions, we started by checking the level of H2BK20ac enrichment in CRISPRi-analyzed enhancers. H2BK20ac is part of the histone H2B N-terminus multisite lysine acetylation (H2BNTac) signature that preferentially marks candidate active enhancers ^48^. Using H2BK20ac ChIP-seq data ^48^, we identified 20,408 H2BK20ac-positive (H2BK20ac^+^) peaks and ranked them from 1 to 20,408, with the highest-to-lowest H2BK20ac enrichment (± 1kb from peak center). 78.8% of all CRISPRi targeted enhancers and 93.3% of high-confidence enhancers overlapped with H2BK20ac^+^ peaks (**Supplemental Table S1A**). Enhancers mapping in top 10,000 strongest H2BK20ac peaks are considered ^high^H2BK20ac enhancers, and those with >10,000 or lacking detectable H2BK20ac are termed ^low,no^H2BK20ac enhancers.

CRISPRi targeted enhancers span the full H2BK20ac range (**Figure 2A**), but those ranked in top 1,000 were selected ∼3 times more often than expected, whereas those ranked >10,000 were selected ∼3 times less frequently than expected (**Figure 2B**). With decreasing H2BK20ac enrichment, the proportion of significant-scoring enhancers decreases (**Figure 2C**, **Supplemental Figure 1B**). In the top 500 ranked enhancers, 34.1% are significant, while merely 3.3-5.9% of ^low,no^H2BK20ac enhancers are significant. Significant scoring enhancers have higher H2BK20ac than non-significant enhancers (p= 6.0e-55, Kolmogorov-Smirnov test) (**Figure 2D**). Of high-confidence enhancers, 61.2% ranked in the top 2,500, 77.6% in top 5,000, and 87.3% in top 10,000. These results show that the level of H2BK20ac enrichment in enhancers strongly predicts functional enhancers.

**Figure 2.**
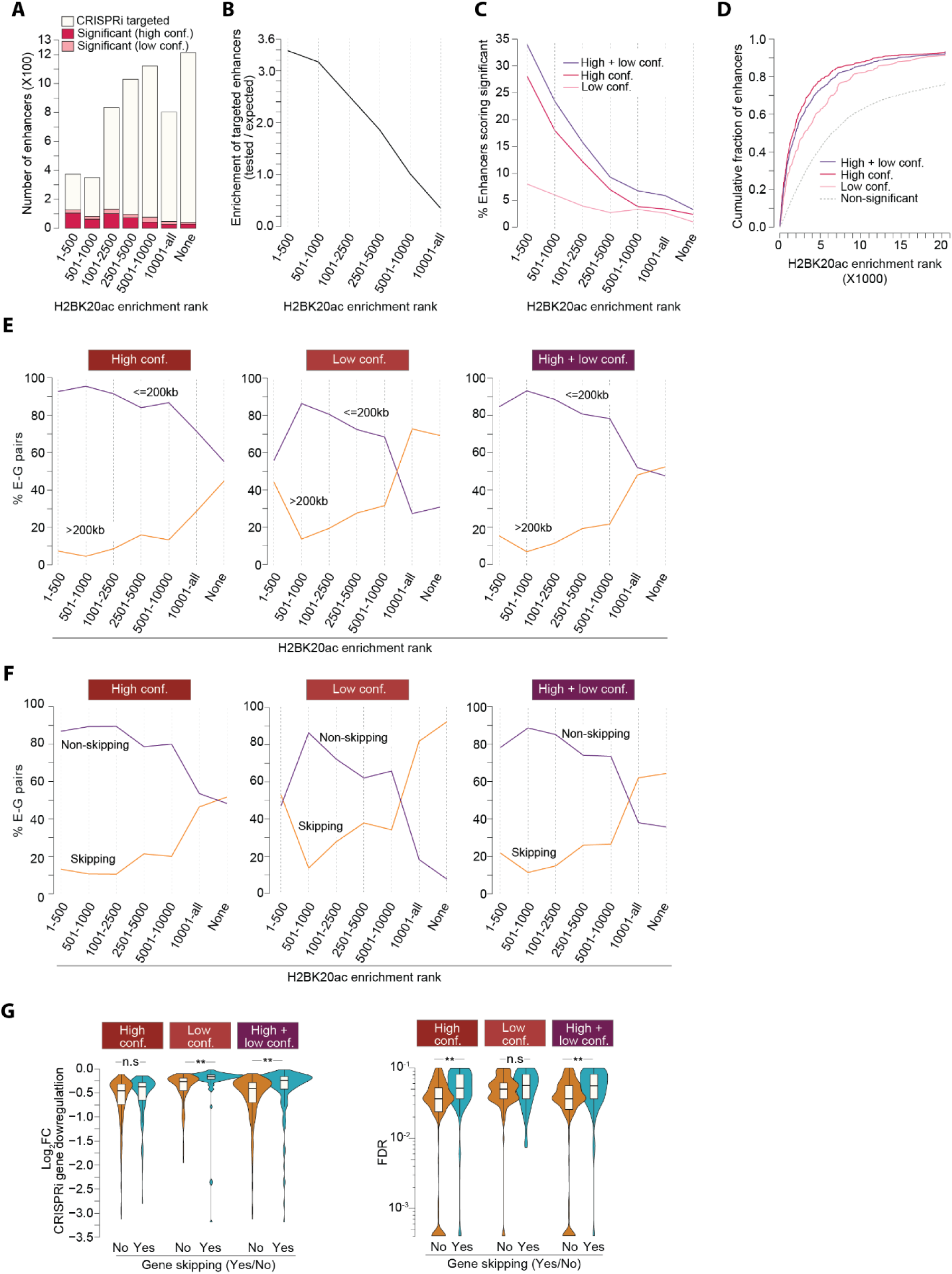
Functional enhancers are marked with strong H2BK20ac, and they impact proximal genes. (**A**) Enhancers are ranked by H2BK20ac ChIP-seq signal within +/- 1kb of the targeted region (see Methods) and grouped into the indicated rank categories. The number of targeted and significant scoring enhancers is shown. High confidence (high conf.) and low confidence (low conf.) enhancer-gene (E-G) pair groups are defined by Gasperini et al.^40^. (**B**) Shown is the relative representation (actual tested/expected) of H2BK20ac marked enhancers among the CRISPRi targeted enhancers. Of note, high H2BK20ac marked are overrepresented, while low H2BK20ac marked enhancers are underrepresented. (**C**) Fraction of CRISPRi significant scoring enhancers across indicated H2BK20ac rank categories. With decreasing H2BK20ac enrichment, the fraction of significant scoring enhancers decreases. (**D**) Cumulative fraction of H2BK20ac-marked enhancers within specified CRISPRi-defined enhancer groups. Non-significant enhancers encompass all CRISPRi-targeted enhancers that did not result in significant gene downregulation. (**E**) Fraction of E-G pairs involving genomic distance of ≤200kb or >200kb in the indicated enhancer H2BK20ac enrichment rank categories and enhancer confidence groups. (**F**) Fraction of active gene-skipping and non-skipping E-G pairs in the indicated enhancer H2BK20ac enrichment rank categories and enhancer confidence groups. (**G**) CRISPRi-induced fold downregulation of enhancer target genes (right panels), and false discovery rate (FDR, left panels), within the specified enhancer confidence groups. Two-sided Mann–Whitney U-test, adjusted for multiple comparisons with the Benjamini–Hochberg method; n.s not significant, \**P*< 0.05, \*\**P*<1e-7.

### H2BK20ac-positive enhancers activate proximal genes

Next, we examined H2BK20ac enrichment in enhancers activating short-range and long-range targets, defined using an arbitrary E-G pair distance threshold of ≤200kb or >200kb. Strikingly, in the top 2,500 H2BK20ac rank category, only 7.2% of ^highconf^E-G pairs have a distance >200kb (**Figure 2E**). In contrast, in the ^low,no^H2BK20ac category, 28.6-44.8% of ^highconf^E-G pairs and 69.2-72.7% of ^lowconf^E-G pairs have >200kb distance.

The propensity of gene-skipping is related inversely to H2BK20ac enrichment in enhancers. In the high-confidence group, only 11.6% of the top 2,500 ranked E-G pairs are skipping, whereas 49.1% of ^low,no^H2BK20ac E-G pairs are skipping (**Figure 2F**). In the low-confidence group, a staggering 85% of ^low,no^H2BK20ac E-G pairs are gene-skipping.

Gene-skipping enhancers cause significantly weaker gene regulation and have a higher false discovery rate (FDR) than non-skipping enhancers (**Figure 2G**). Furthermore, targets of non-skipping enhancers are biased for cell-type-specific genes, while gene-skipping enhancers show no such preference (non-skipping genes versus expressed genes, p=2.2e-8; skipping genes versus expressed genes, p=0.26; Kolmogorov-Smirnov test) (**Supplemental Figure 1C**).

We also checked H2BK20ac enrichment in enhancers activating single or multiple genes, proximal or distal genes, or combinations thereof (**Supplemental Figure 1D**). 79.5% of high-confidence enhancers activated a single proximal gene, while 15% activated a single distal gene. Only 5.5% (24/441) of high-confidence enhancers activated multiple genes and over half of them activated consecutively located proximal genes without avoiding the activation of intervening genes.

These results reveal that gene-skipping and non-skipping enhancers differ systematically in H2BK20ac abundance, gene activation ability, FDR, and cell-type-specificity of their target genes.

### H2BK20ac-positive enhancers activate genes using CBP/p300

Next, we investigated the requirement of CBP/p300, the enzyme catalyzing H2BNTac^49^, for enhancer function. CBP/p300-regulated genes have been quantified in K562 ^50^ using the CBP/p300 catalytic inhibitor A-485 ^51^. For 94% of CRISPRi-defined enhancer targets, CBP/p300-dependent transcription regulation data were available (**Supplemental Fig 2A**). A-485-regulated genes were categorized as highly downregulated (HD, ≥2-fold down), intermediate downregulated (ID, ≥1.5 fold down), slightly downregulated (≥1.2-fold down, SD), not downregulated (ND, <1.2-fold down), or not quantified (NA). The fraction of regulated genes in K562 was as follows: 9.5% HD, 7.0% ID, 11.2% SD, and 72.3% ND. If enhancer targets were not biased for regulation by CBP/p300, then by random, we would anticipate the same proportion of A-485 regulated genes among CRISPRi-identified enhancer targets.

Notably, in ^highconf^E-G pairs, 40.6% include HD genes (HD versus ND gene group, odds ratio 9.9, adjusted p-value 7.7e-79, fisher test) (**Figure 3A**). The requirement of CBP/p300 scales with H2BK20ac enrichment; as enhancer H2BK20ac levels decrease, the proportion of HD+ID genes decreases while ND+NA genes increase (**Figure 3B, Supplemental Figure 2B**). In the high-confidence group, in top 1,000 H2BK20ac enrichment rank, only 18.1% E-G pairs include ND+NA genes, whereas, in the ^low,no^H2BK20ac rank, 75.4% E-G pairs include ND+NA genes (**Figure 3B**). These results demonstrate that CBP/p300 is preferentially required for activating the targets of H2BNTac-positive enhancers.

**Figure 3.**
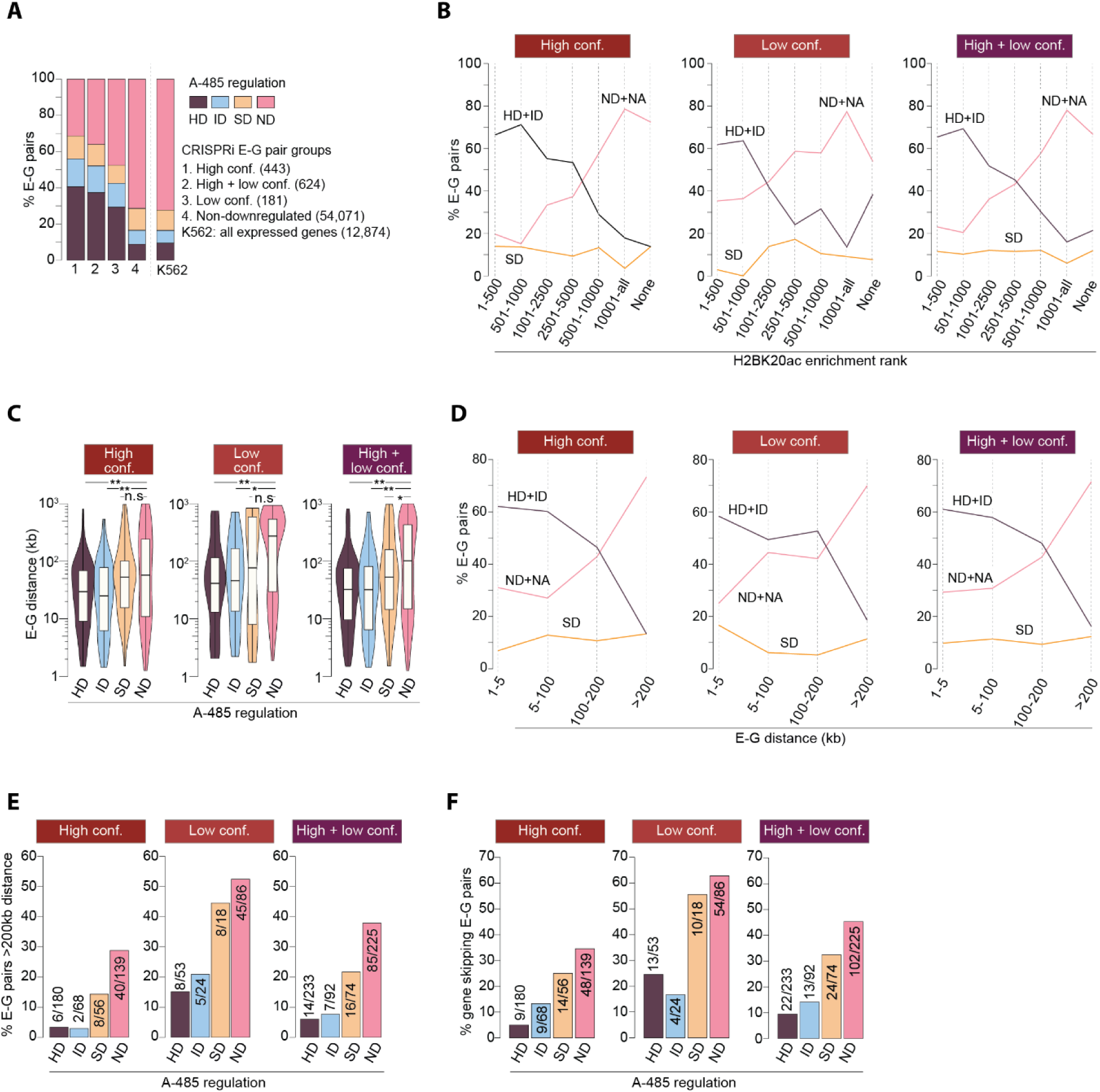
Gene-skipping and non-skipping enhancers distinctly require CBP/p300. (**A**) Fraction of genes regulated by A-485 within the indicated groups of E-G pairs. A-485-induced regulation of K562-expressed genes is shown as a reference. Genes are classified based on A-485-induced nascent transcript changes as highly downregulated (HD, ≥2-fold down), intermediate downregulated (ID, ≥1.5-fold down), slightly downregulated (SD, ≥1.2-fold down), or not downregulated (ND, <1.2-fold down). (**B**) Fraction of E-G pairs involving the indicated A-485-regulated gene class in the specified enhancer H2BK20ac enrichment rank categories and enhancer confidence groups. A-485 regulated genes are classified as described in Figure 2A. NA: A-485-induced gene regulation data not available. (**C**) E-G pairs are categorized according to A-485-induced regulation of enhancer targets in the indicated enhancer confidence groups. E-G pair genomic distances are plotted within the indicated A-485 gene regulation categories. Two-sided Mann–Whitney U-test, adjusted for multiple comparisons with the Benjamini–Hochberg method; n.s not significant, \**P*< 0.05, \*\**P*<0.01. (**D**) Fraction of E-G pairs involving the indicated A-485-regulated gene class in the specified E-G pair distance categories and enhancer confidence groups. A-485 regulated genes are classified as described in Figure 2A. NA: A-485-induced gene regulation data not available. (**E-F**) The fraction of enhancers that activate target genes located >200 kb away (E) or skip proximal active genes (F) within the specified A-485-regulated gene classes and indicated enhancer confidence groups.

### CBP/p300-dependent enhancers seldom skip active genes

In the high-confidence group, E-G pair distance is twice as long in the ND category than in the HD+ID category (median 28.7kb versus 57.5kb, p= 8.2e-5, Mann-Whitney U-test), and E-G pair distances are even longer in the low confidence group (**Figure 3C**). The low-confidence group includes ∼2.8 times more E-G pairs with >200kb distance than the high-confidence group (36.1% versus 12.8%, odds ratio 3.9, p= 3.9e-11, fisher test) (**Supplemental Figure 2C**). Most E-G pairs with <100kb distance involve HD+ID genes, whereas those with >200kb distances mainly involve ND genes (**Figure 3D, Supplemental Figure 2D**). In the high-confidence group, 60.5% of E-G pairs with distances <100kb feature HD+ID genes. In contrast, in E-G pairs with a >200kb distance, the fraction of HD+ID genes is the same as expected by random by chance (**Figure 3D**, **Figure 3A**). In this group, the prevalence of E-G pairs with distances >200kb is 9 times more in the ND group than in the HD+ID group (3.2% versus 28.8%, odds ratio 12.0, p= 7.0e-13, fisher test) (**Figure 3E**). This finding shows that a mere 3.2% (8/248) of CBP/p300-dependent high-confidence enhancers activate genes located >200kb away.

The propensity of gene-skipping increases as the dependency of genes on CBP/p300 weakens (**Figure 3F**). In the low-confidence group, 62.8% of ND category E-G pairs involve skipping. In contrast, in the high-confidence group, only 7.3% of HD+ID category E-G pairs involve skipping. These computational analyses show that 7.3% (18/248) of CBP/p300-dependent high-confidence enhancers skip active genes. Still, manual examination of these enhancers indicates that they most likely activate their proximal active genes rather than skipping them (**Supplemental Note 1**, **Supplemental Table S1B**).

Prompted by this observation, all 664 significant scoring E-G pairs were examined manually. According to manual classification, 64.3% of E-G pairs are confirmed as likely/plausible targets of CRISPRi-identified enhancers that function using CBP/p300 (Group 1), 26.7% are deemed either false positives, indirect enhancer targets, or targets of enhancers that function without CBP/p300 or (Group 2), while 9% of E-G pairs could not be classified due to various reasons (Group 3) (**Supplemental Note 2**, **Supplemental Table S1C**). Most Group 1 enhancers regulate a single gene, and few regulate 2 to 3 genes, either directly flanking the enhancer or located consecutively on one side of the enhancer. In Group 1, we only found 3 E-G pairs where enhancers may entirely skip active gene(s) to activate a distal target exclusively. These analyses indicate that CBP/p300-dependent, H2BK20ac-marked high-confidence enhancers predominantly activate proximal genes and seldom (∼1-2%) skip genes, whereas most CBP/p300-independent enhancers skip active genes.

While skipping of active genes is seldom, in many instances, enhancers do activate non-neighboring genes, instead of the nearest neighbor (**Figure 4**). For example, enhancers activating *ZNF518B*, *NR2F2*, and *TFRC* do not affect their nearest active genes. In the CRISPRi data, some enhancers, for example those near *FGF2*, *PICALM*, and *SLC6A6*, are connected to both nearest and non-nearest neighbor genes, but the nearest genes show stronger downregulation by CRISPRi. Consistent with CRISPRi data, in all these instances, CBP/p300 inhibition shows that enhancers mainly regulate the non-nearest neighbor genes. This shows that in these instances, E-P proximity alone is not sufficient to predict which of the neighboring genes are regulated by enhancers, but combination of the enhancer proximity and the dependency of proximal genes on CBP/p300 can help identify genes that are most likely gene affected by enhancers.

**Figure 4.**
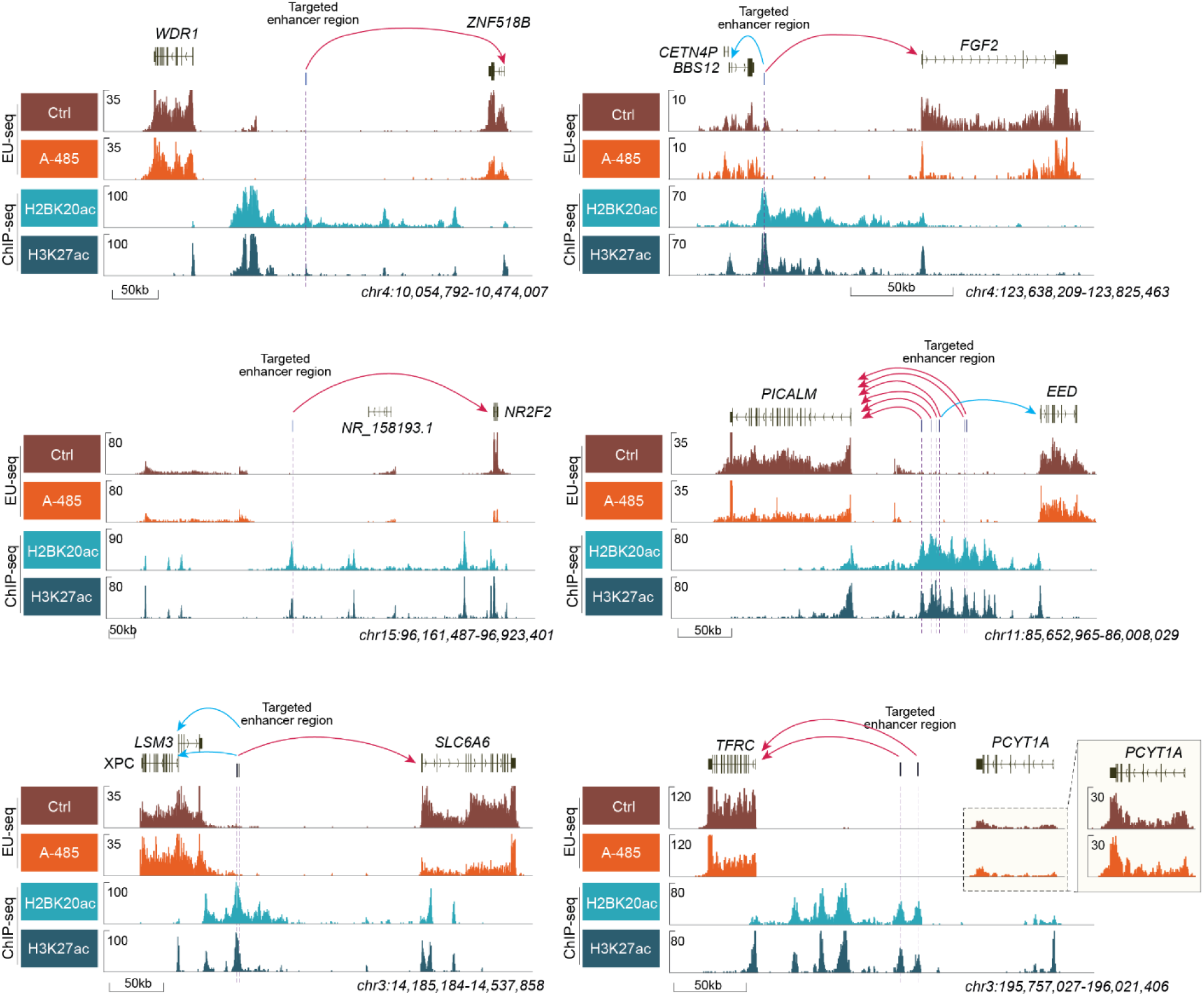
Some enhancers activate non-nearest neighbor active genes at long genomic distances. Representative genome browser tracks of CRISPRi-defined E-G pairs involving enhancers that activate non-nearest neighbor genes or are linked to multiple genes. In three instances (*ZNF518B*, *NR2F2*, and *TFRC*), one of the neighboring genes is defined as an enhancer target by CRISPRi. Only the CRISPRi-defined enhancer target is downregulated between the enhancer neighbor genes by acute CBP/p300 inhibition. In the remaining three instances (*FGF2*, *PICALM*, and *SLC6A6*), some enhancers are linked to more than one gene. In each of these cases, the genes that are most strongly downregulated by CRISPRi (*FGF2*, *PICALM*, and *SLC6A6*) are more prominently downregulated by CBP/p300 inhibition, whereas genes that are weakly downregulated by CRISPRi either show very weak downregulation (*EED*), or no appreciable downregulation (*BBS12*, *LSM3*, and *XPC*) after CBP/p300 inhibition. These examples confirm that enhancers can activate genes at long genomic distances and preferentially impact non-nearest neighbor genes, but they tend to activate one of the neighboring genes preferentially. All genes, except for *EED*, are identified as high-confidence enhancer targets by CRISPRi. *EED* is identified as a low-confidence target. For enhancers linked to more than one gene, red arrows show genes that CRISPRi most strongly downregulates, while blue arrows show genes that are downregulated relatively weakly by CRISPRi.

### CBP/p300 is necessary for open but not closed chromatin enhancers

In massively parallel reporter assay (MPRA), distinct human enhancers activate genes through different coactivators ^52^. Prompted by the above findings, we examined the properties of enhancers functioning with or without CBP/p300 in MPRA. Remarkably, among the examined coactivators (BRD2, BRD4, MED14, CDK7, CDK9, CBP/p300), CBP/p300 is the only coactivator whose requirement strongly aligns with both H2BK20ac enrichment in enhancers and their accessibility in native chromatin (**Supplemental Figure 3A-B**). High H2BK20ac marked enhancers require CBP/p300 and exhibit open accessibility, while enhancers functioning without CBP/p300 show low or no H2BK20ac and poor accessibility. This result indicates that enhancers with open chromatin accessibility and high H2BK20ac require CBP/p300, while those with closed chromatin accessibility and low or no H2BK20ac do not. Enhancers generally become active after gaining chromatin accessibility ^3^. Nonetheless, closed enhancers can activate genes in plasmid-based reporters ^53,54^ and may do so in native chromatin ^55^. Our results imply that they possibly utilize coactivators other than CBP/p300 if they do.

### An independent CRISPRi dataset validates enhancer attributes

In a focused effort, Morris et al. CRISPRi targeted 543 blood trait GWAS-harboring candidate enhancers and defined 135 as functional ^41^. Of these 135 GWASs, CRISPRi targeting led to gene downregulation in 125, upregulation in 7, and up- and down-regulation in 3 cases. The targeted GWAS regions overlapped minimally with enhancers targeted by Gasperini et al. (**Figure 5A**). Thus, the dataset provided a valuable resource for validating and expanding the above findings.

**Figure 5.**
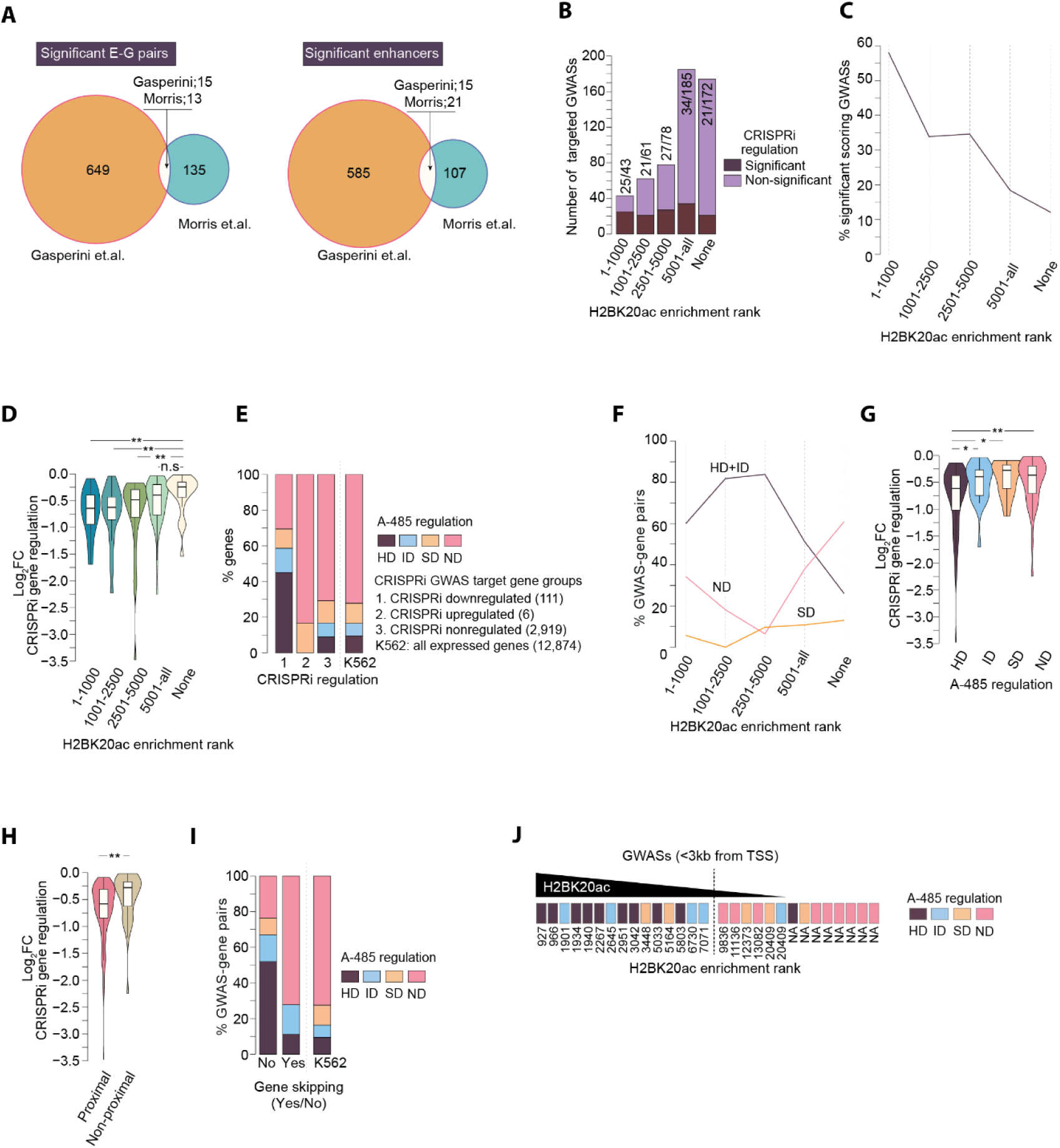
CRISPRi-targeted non-coding GWASs independently validate key enhancer attributes identified in the genome-scale CRISPRi dataset. (**A**) The overlap between significant scoring enhancers and GWASs (right panel) and E-G pairs and GWAS-gene pairs (left panel) in the Gasperini et al. and Morris et al. datasets^40,41^. Enhancers and GWASs are considered overlapping if they are positioned within a 1kb distance. Only the CRISPRi downregulated GWAS-gene pairs from the Morris et al. dataset are included in this analysis. (**B**) CRISPRi-targeted GWAS loci are ranked based on H2BK20ac enrichment (see Methods) and grouped into the specified H2BK20ac rank categories. The number of CRISPRi-targeted GWAS loci and significant scoring GWASs are shown. "None" refers to GWASs without detectable H2BK20ac. (**C**) Fraction of significant scoring GWASs within the indicated H2BK20ac rank categories. (**D**) CRISPRi-induced downregulation of GWAS target genes in the indicated H2BK20ac rank categories. Two-sided Mann–Whitney U-test, adjusted for multiple comparisons with the Benjamini–Hochberg method; n.s not significant, \*\**P*<0.01. (**E**) Fraction of A-485 regulated gene class within the indicated categories of GWAS target genes. GWAS target genes are categorized based on their regulation by CRISPRi: “CRISPRi downregulated” refers to genes significantly downregulated by CRISPRi; “CRISPRi upregulated” refers to genes significantly upregulated by CRISPRi; “CRIPSRi nonregulated” indicates genes whose expression is not downregulated by CRISPRi (log2 FC > -log_2_ (1.05) and *P*>0.05 in CRISPRi experiment, and genes are not involved in any significant E-G pairs). (**F**) Fraction of GWAS-gene pairs involving the indicated A-485 regulated gene class within the specified GWAS H2BK20ac enrichment rank categories. (**G-H**) CRISPRi-induced downregulation of GWAS target genes in the indicated A-485-regulate gene class (F), or gene proximity class (G). GWAS targets are classified based on the regulation of GWAS target genes by A-485 (F), or based on whether GWAS is connected to the proximal (nearest) or non-proximal (non-nearest) gene (G). Two-sided Mann– Whitney U-test, adjusted for multiple comparisons with the Benjamini–Hochberg method; \**P*<0.05, \*\**P*<0.01. (**I**) The fraction of A-485 regulated genes in the skipping ("yes") and non-skipping ("no") GWAS groups. As a reference, the fraction of A-485-regulated genes in K562 is shown. (**J**) Shown is A-485-induced regulation of significant scoring GWAS targets with GWAS-target gene TSS distance of <3kb. GWASs are rank ordered based on H2BK20ac enrichment, and the dotted line demarcates GWASs with high and low H2BK20ac enrichment.

About half of the analyzed GWASs had ^low,no^H2BK20ac (H2BK20ac rank >10,000: 19.1%; no detectable H2BK20ac: 31.9%) (**Supplemental Table S2A**). Of the GWASs ranking top 1,000 H2BK20ac peaks, 58.1% scored as significant, whereas for those with no H2BK20ac, only 12.2% scored significant (**Figure 5B-C**). GWASs with strong H2BK20ac caused significantly stronger gene downregulation in CRISPRi (adjusted p-value: top 1000, 2.2e-3; top 1001-2500, 6.7e-3; top 2501-5000 6.7e-3, each compared with ^no^H2BK20ac, Mann-Whitney U-test) (**Figure 5D**).

The GWAS targets showing decreased expression by CRISPRi silencing were biased for A-485-induced downregulation, whereas those showing increased expression were not (**Figure 5E**). With decreasing H2BK20ac enrichment in GWASs, the proportion of HD+ID genes diminished among GWAS targets (**Figure 5F**). Notably, GWASs linked to ND genes cause weaker gene regulation in CRISPRi than those linked to HD+ID genes (**Figure 5G**), and GWAS linked to non-proximal genes cause weaker gene downregulation than those linked to proximal genes (**Figure 5H**). Non-skipping GWAS targets are prominently downregulated by A-485 (HD+ID versus ND group, odds ratio 7.2, p= 2.3e-4, fisher test), whereas gene-skipping GWAS targets show no bias for A-485-induced gene regulation (HD+ID versus ND group, odds ratio 1.68, p= 0.36, fisher test) (**Figure 5I**).

Interestingly, H2BK20ac enrichment in GWASs can reliably predict the strength of individual GWASs, except for GWASs that cause very weak gene downregulation by CRISPRi, or the targeted GWASs are present within 3kb of the TSS their target genes (see **Supplemental Note 3**, **Supplemental Figure 4A-C**). 19.6% of significant scoring GWASs are present within 3kb of their target gene TSS. We analyzed H2BK20ac and A-485-induced regulation of GWAS-gene pairs with <3kb distance. Half of these GWASs were marked with high H2BK20ac, and the genes linked to them were downregulated by A-485 (9 HD, 4 ID, 2 SD). The remaining half had weak or no H2BK20ac, and the genes linked to them were minimally impacted by A-485 (1 HD, 1 ID, 3 SD, 9 ND) (**Figure 4J**). This implies that CRISPRi possibly overestimates the function of promoter-proximal GWASs by directly repressing their promoters.

All significant GWAS-gene pairs were examined manually. Among H2BK20ac^+^ functional GWASs, only two GWASs (rs2238368, rs11864973) were confirmed to skip active genes, and both skip the proximal NPRL3 to activate alpha-globin genes. These analyses confirm that H2BNTac enrichment predicts enhancer function, and CBP/p300-dependent enhancers seldom skip active genes.

We also checked why some H2BK20ac-positive GWASs score significantly in CRISPRi while others don’t. Our analyses suggest that many of the non-significant scoring H2BK20ac^+^ GWASs are likely functional, but they are located distally from their putative targets, and their quantitative impact is subtle compared to significant scoring GWASs (see **Supplemental Note 4**, **Supplemental Figure 5A-H**). This implies that CRISPRi can identify targets of strong GWASs but is likely underpowered to detect targets of weak GWASs, which cause weaker changes in transcription. These results demonstrate that H2BK20ac-positive GWASs activate genes using CBP/p300 and seldom skip active genes.

### CBP/p300-dependent enhancers operate within a limited distance range

Recently, it has been proposed that interaction between enhancer RNA (eRNA) and promoter upstream transcripts (PROMPTs) dictate E-P promoter specificity, and eRNA knockdown of selected enhancers showed that enhancers frequently activate genes located >1Mb ^28^. This would suggest that restricting the enhancer target search window to 500kb to 1Mb in CRISPRi analyses may cause systematic underestimation of distal targets. We reanalyzed the Gasperini et al. dataset using different target search windows (ranging from 30kb to 20Mb) and FDRs (0.01, 0.05, and 0.1) to examine how search parameters influence the nature of identified targets.

At FDRs of 0.05 and 0.1, the identified E-G pairs peaked around 200kb, then steadily decreased (**Figure 6A, Supplemental Figure 6A**). Restricting the search window to 200kb led to identifying 27.3% more E-G pairs than a 1Mb window at FDR 0.1 and 58.8% more at FDR 0.05. At FDR 0.01, significant E-P pairs peaked at ∼100kb, dropping their numbers nearly four-fold with extending search windows to ≥500kb.

**Figure 6.**
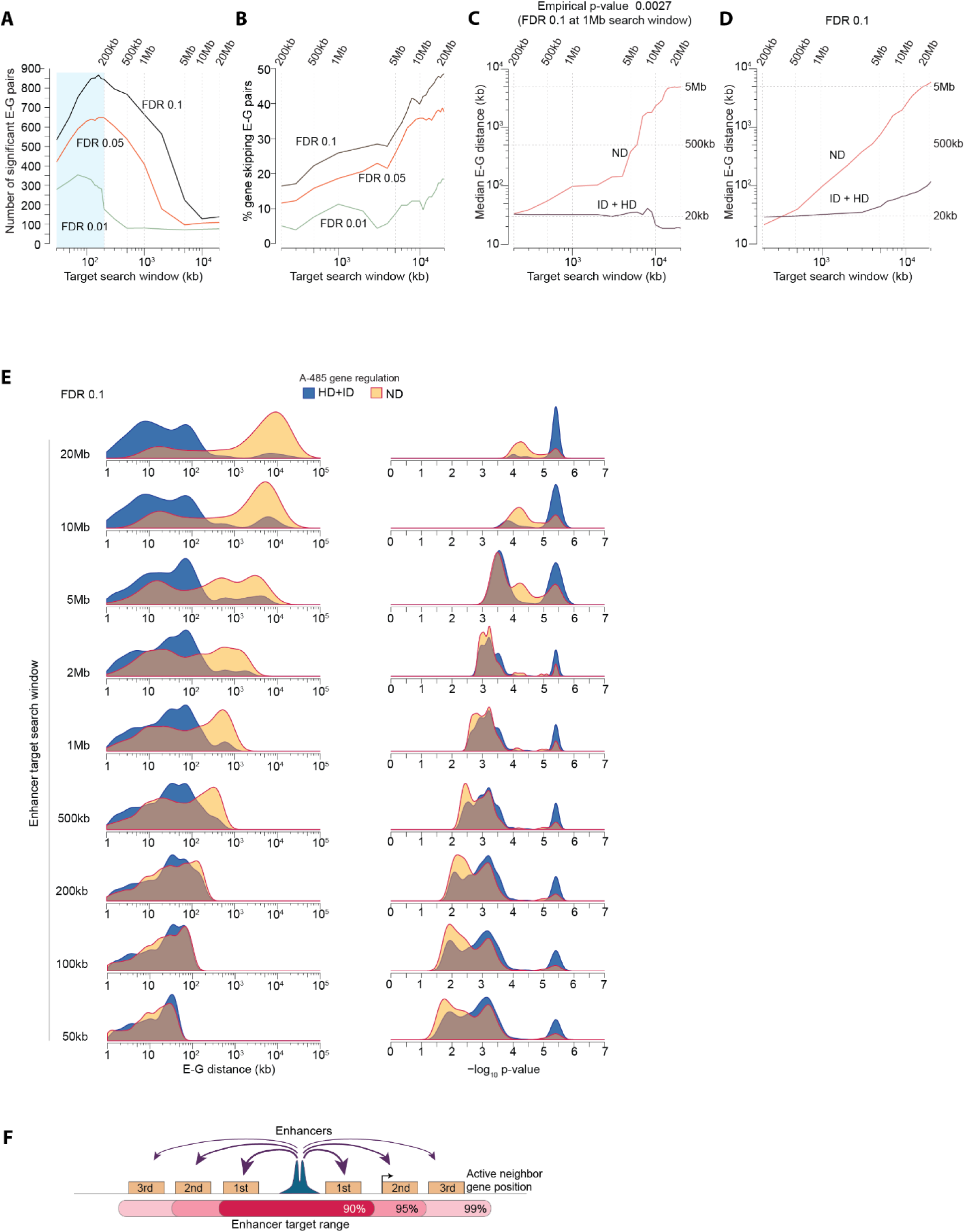
H2BK20ac-marked enhancers act within a limited distance range. (**A**) Number of significant scoring E-G pairs at the specified false discovery rates (FDRs) and indicated enhancer target search windows. E-G pairs were searched in the Gasperini et al. dataset^40^ by varying the target search window from +/- 30kb to 20Mb. (**B**) Fraction of gene skipping E-G pairs identified at the indicated FDRs and enhancer target search window. E-G pairs were identified by varying FDRs and target search windows, as specified in Figure 4A. (**C-D**) The median E-G pair distance for the specified groups of A-485-regulated gene class in the indicated FDRs and target search windows. Enhancer targets were searched at target search windows varying from +/-30kb to 20Mb. Significant scoring E-G pairs were identified by applying a fixed FDR (0.1) (C), or by using a fixed nominal p-value (p = 0.0027, p-value fixed to yield FDR of 0.1 at 1Mb) (D). (**E**) Shown are genomic distances of E-G pairs involving A-485 downregulated (HD+ID) and not downregulated (ND) genes (left panels), as well as nominal p-values of the identified E-G pairs (right panels). E-G pairs are identified using the indicated genomic distance windows and FDR of 0.1. (**F**) Proposed range of H2BK20ac+ enhancer target genes.

While the number of significant E-P pairs decreases with increasing distance and higher FDR, the proportion of gene-skipping E-G pairs increases (**Figure 6B**). Using the same parameters (FDR 0.05, distance window of 500kb), Gasperini et al. and Morris et al. datasets showed similar proportions of gene-skipping E-G pairs (15.8% versus 13.5%), indicating that differences in FDR and distance thresholds account for the two-fold difference in frequencies of gene-skipping enhancers in these datasets.^40,41^.

Upon noticing this trend, we investigated E-G pairs identified with fixed FDR (variable p-values) or fixed nominal p-value (variable FDR). At a fixed FDR, widening the search window increased the E-G pair distance, but only for pairs involving ND genes (**Figure 6C, Supplemental Figure 6B**). Indeed, the median E-G pair distance increased by over two orders of magnitude in the ND group but remained almost unchanged in the HD+ID group (**Figure 6C**).

At a fixed empirical p-value cutoff (p= 0.0027, corresponding to FDR 0.1 at the 1Mb window), broadening the search window resulted in a gradual rise in the proportion of enhancers linked to ND genes (**Supplemental Figure 6C**), along with a disproportionate increase in E-G pair distance in the ND group compared to the HD+ID group (**Supplemental Figure 6D**). Notably, the median E-G pair distance in the ND group exhibits a linear increase in the log-log plot, indicating a power-law relationship (**Figure 6D**). Additionally, E-G pairs involving ND genes tended to have lower statistical confidence across all distances (**Figure 6E**).

These results demonstrate that the selection of the target search window and FDR not only impacts the number of significant scoring E-G pairs, but also systematically alters the nature of identified enhancer target genes. Thus, the choice of target search window directly influences the conclusions about the proportion of gene-skipping enhancers and the dependency of enhancers on CBP/p300.

### Properties of RIC-seq-identified enhancers do not conform with those identified by CRISPRi

The above results prompted us to examine the properties of E-P pairs identified in RIC-seq, which suggests that the complementarity between enhancer RNA and promoter upstream transcripts dictates E-P specificity ^28^. In K562, RIC-seq linked 3,091 enhancers to 3,780 promoter groups, forming 7,030 E-P pairs, as many promoter groups included multiple promoters (**Supplemental Table S4**). Most enhancers are associated with distal genes, with a staggering 78% forming inter-chromosome E-P contacts (**Figure 7A**). Over 70% of E-P pairs involved enhancers interacting with multiple promoters (**Figure 7B**).

**Figure 7.**
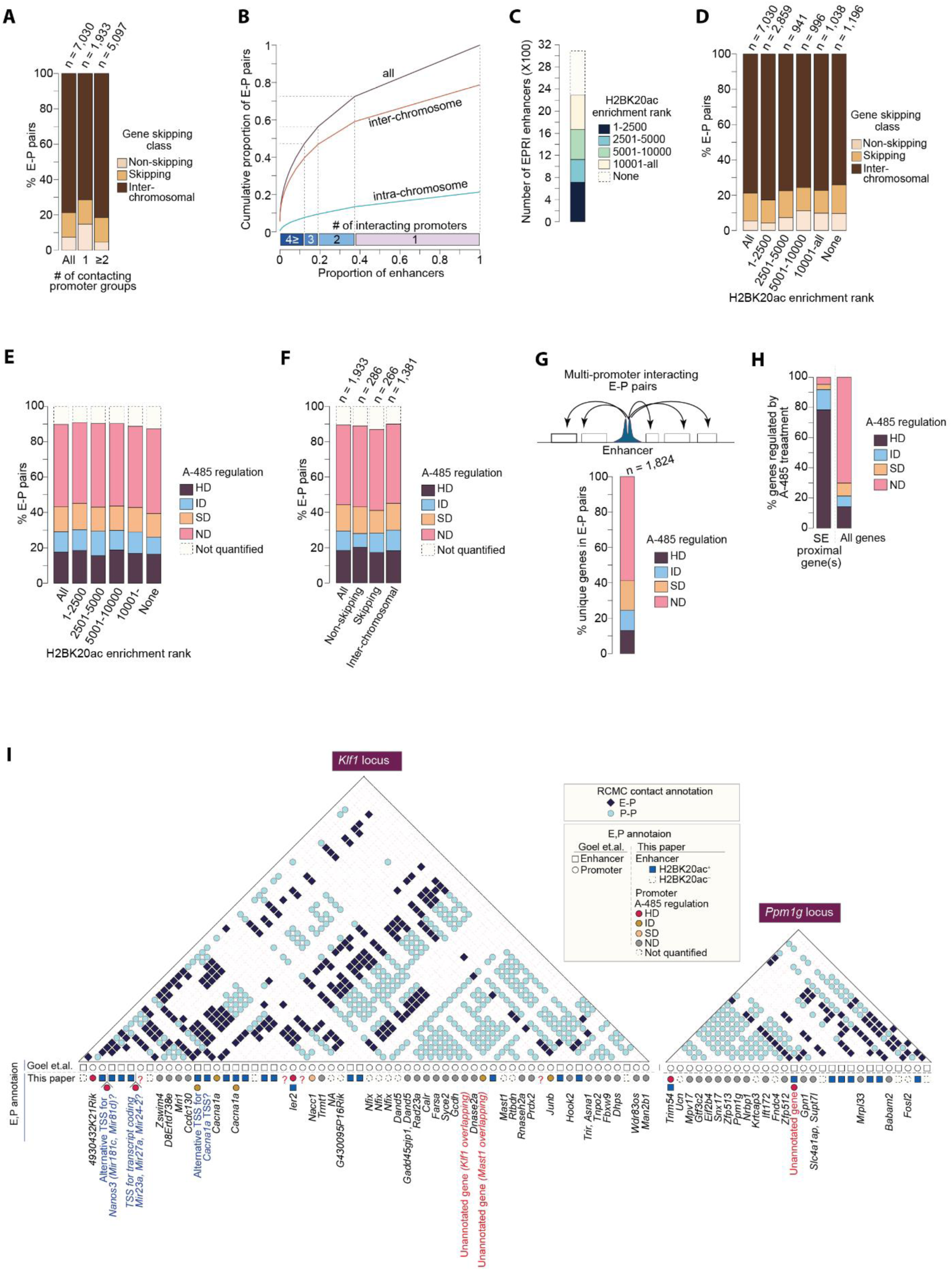
MED1 enrichment-defined super-enhancers share properties with CRISPRi-identified enhancers, but multi-way interacting promoters in RIC-seq and RIC-seq exhibit distinct regulation by CBP/p300. (**A**) Number of RIC-seq-identified E-P pairs in K562 cells^28^. Enhancers are classified as follows: Non-skipping enhancers contact proximal active genes; skipping: enhancers contact non-proximal active genes; inter-chromosomal: enhancers contact target genes occurring in chromosomes different from those harboring the enhancers. Enhancers linked to single (1) and multiple (≥2) promoter groups are analyzed separately. (**B**) Proportion of enhancers engaged in 1, 2, 3, or ≥4 E-P contacts. The proportion of E-P pairs involving inter-chromosome, intra-chromosome, and all contacts are displayed. (**C**) The number of RIC-seq-identified candidate enhancers overlapping with H2BK20ac-positive peaks within the specified H2BK20ac enrichment rank categories. (**D**) Fraction of E-P pairs involving non-skipping, skipping, and inter-chromosomal target genes within the indicated groups of enhancer H2BK20ac enrichment rank categories. (**E**) Fraction of E-P pairs involving the indicated class of A-485 regulated genes within the specified groups of enhancer H2BK20ac enrichment rank categories. (**F**) Fraction of E-P pairs involving the indicated class of A-485 regulated genes within non-skipping, skipping, and inter-chromosomal RIC-seq E-P pair groups. A-485-induced gene regulation is analyzed for RIC-seq-identified enhancers that are exclusively linked to a single promoter group. (**G**) A-485-induced regulation of the genes contacting putative CBP/p300-dependent, multi-promoter linked RIC-seq enhancers. Among multi-promoters contacting RIC-seq enhancers, putative CBP/p300-dependent enhancers were identified as having at least one gene downregulated by A-485 by ≥2-fold. After excluding the strongest A-485-regulated gene from each enhancer, regulation of the remaining genes by A-485 is depicted. Some enhancers are linked to overlapping promoter groups. This analysis only includes unique genes occurring in the promoter groups to avoid counting the same genes multiple times. (**H**) A-485-induced regulation of immediate neighbor gene(s) of MED1-enrichment-defined SEs (n= 231) ^43^. A-485-induced regulation of all mESC genes is shown as a reference. (**I**) RCMC detected interactions between candidate enhancers and promoters and regulation of genes after CBP/p300 inhibition. The E-P and P-P contacts are shown based on original annotations by Goel et al. ^34^. Original annotations are shown. We also re-annotated candidate enhancers and promoters. Candidate active enhancers are identified by H2BK20ac, active promoters by H3K4me3, and active genes by nascent transcript expression in mESC. A few of the elements, such as the promoter of *Ier2*, are marked with both strong H2BK20ac and H3K4me3, and thus, they are annotated as both enhancer and promoter. Expressed genes, as well as genes regulated by CBP/p300 inhibition, are shown. Putative unannotated transcripts, and alternative transcripts of known genes are indicated. To minimize crowding, contacts involving E-E interaction and interaction with CTCF peaks are not shown.

About half of RIC-seq enhancers rank among the strongest 10,000 H2BK20ac peaks (**Figure 7C**). Surprisingly, unlike CRISPRi-identified enhancers (**Figure 2F**), H2BK20ac enrichment in RIC-seq-identified enhancers showed no association with the extent of gene skipping (**Figure 7D**), or with A-485-induced regulation of enhancer-interacting promoters (**Figure 7E**). The fraction of A-485 regulated genes did not differ among skipping, non-skipping, and inter-chromosomal enhancer targets (**Figure 7F**).

Functional analyses of five RIC-seq-identified enhancers showed that each regulates multiple genes, some locating tens of megabases away from enhancers ^28^. We posited that the same enhancers likely activate their targets using the same coactivators. We examined enhancers that interact with multiple promoter groups and where at least one of the interacting genes is highly downregulated (>2-fold) by A-485. Because these enhancers include at least one gene regulated by A-485, we presume they function using CBP/p300. To examine the co-activation of other interacting genes by these enhancers, for each enhancer, we excluded the most strongly affected A-485 downregulated gene based on which these enhancers were identified. Most of the remaining genes linked to these enhancers are not downregulated of A-485 (**Figure 7G**). This shows that promoters interacting with the same enhancers in RIC-seq are activated by different coactivators.

These results uncover fundamental differences between E-P pairs identified by RIC-seq and CRISPRi. While the cause of this inconsistency remains unclear, we can only envision two scenarios. (1) RIC-seq detects unique functional E-P relationships not captured by CRISPRi, or (2) RIC-seq-identified E-P contacts poorly inform about true enhancer targets. We consider the latter possibility more likely. We note that many studies reporting a high prevalence of inter-chromosomal E-P contacts have used formaldehyde as a single fixative ^15,28–31^. Inter-chromosome contacts in HiC and Micro-C decrease many fold when cells are fixed with double fixatives compared to formaldehyde alone ^56^, leading some studies to use inter-chromosome interactions to estimate background noise ^44^. Regardless of the reasons, our results show a stark contrast in the properties of RIC-seq and CRISPRi-identified enhancers.

### Properties of MED1-enriched super-enhancers conform to those of CRISPRi-defined enhancers

To check whether enhancers defined based MED1 enrichment follow the rules of CRISPRi-identified enhancers or RIC-seq-identified enhancers, we examined the properties of 231 SEs defined by high MED1-enrichment in mESC ^43^. SE candidate target genes were inferred by the regulation of proximal genes by CBP/300 inhibition. Over 90% of SEs (211/231) had immediate neighboring gene(s) downregulated by ≥1.5-fold by A-485, including all 12 genes previously identified as SE targets^57^ (**Figure 7H**, **Supplemental Table S5**). Putative targets of SEs included specific gene isoforms and unannotated transcripts.

About 5% of SEs (17/231) showed no indication of regulation of proximal genes (up to 5 genes away from SEs), and if the proximal genes were affected, they had other H2BK20ac-positive proximal enhancers (**Supplemental Table S5**). We found that one of these putative SEs (mSE_00129) is a promoter rather than an enhancer, and several others have weak or no H2BK20ac enrichment (mSE_00164, mSE_00219, mSE_00076), suggesting these SEs have weak or no enhancer activity, or function without using CBP/p300, or regulate distal genes not identified by us. This shows that, like CRISPRi-identified enhancers, most MED1-enriched SEs are marked by H2BK20ac and predominantly impact their proximal genes using CBP/p300.

### Multi-way E-P interactions in microcompartments poorly inform enhancer targets

Finally, we checked the functional significance of recently reported nested E-P microcompartments ^34^. We examined *Klf1* and *Ppm1*g loci in mESC, where candidate enhancers show extensive interactions with proximal and distal genes ^34^. According to our manual annotations, the *Klf1* locus harbors 30 actively transcribed genes, including three annotated ones (**Figure 7I**). Two of the unannotated genes overlap with *Klf1* and *Mast1* but are transcribed from the opposite strand. The *Ppm1g* locus contains 15 actively transcribed genes, including one poorly annotated. Both loci also contain several candidate enhancers marked by H2BK20ac. Of the 45 active genes in these two loci, 10 are downregulated by CBP/p300 inhibition, and strikingly, each of them has proximal H2BK20ac-marked active candidate enhancers. The remaining 35 genes are not regulated by CBP/p300, even though they also extensively interact with candidate enhancers and promoters of the genes regulated by CBP/p300. This shows that although enhancers can interact with multiple promoters in RCMC analyses, their proximal and distal interacting genes are regulated distinctly by CBP/p300. This suggests that the regulatory information in enhancers is not commonly shared among all interacting promoters in microcompartments; instead, it remains private to individual gene(s).

## Discussion

The prevailing view is that enhancers frequently interact with multiple promoters, often skip active genes, and activate genes using distinct coactivators ^10,15–32,34,52,58–60^. This work advances current models by showing that: (1) properties of gene-skipping E-P pairs identified by CRISPRi and physical E-P interaction mapping are different. (2) CRISPRi identified active gene-skipping and non-skipping enhancers exhibit distinct attributes. (3) Most CRISPRi-defined enhancers act proximally, seldom skip active genes, and activate their target genes using CBP/p300. (4) The CRISPRi-identified enhancers functioning without CBP/p300 exhibits atypical attributes and act without any distance constraints. Below, we discuss these findings and their conceptual implications for understanding gene regulation by enhancers and GWAS variants.

### Active gene-skipping and multi-way E-P interactions are poor indicators of functional relationships

Mapping E-P interactions is one of the most widely used approaches for inferring enhancer targets in mammalian cells. The underlying assumption is that 3D proximity is a better predictor of functional E-P relationships than linear E-P distance. Our results challenge this notion. We show a striking contrast between the properties of enhancer targets nominated by RIC-seq and CRISPRi. Furthermore, we reveal systematic differences in the validation rates of gene-skipping and non-skipping E-P pairs identified by Micro-C, with gene-skipping interactions showing poor validation by CRISPRi. Also, short-range (<20 kb) E-P interactions are underdetected in Micro-C, and this issue is likely even more pronounced with methods like HiC and RIC-seq.

This distance limitation can be overcome by mapping interactions at ultra-high resolution by locus-specific methods like Tri-C, MCC, and RCMC ^33,34,61^. However, with deeper mapping, detected interactions become less specific and tend to mirror chromatin accessibility landscape ^34,61^. For example, in the *Ppm1g* and *Klf1* loci, virtually all promoters and enhancers engage in multi-way interaction. If the entire mammalian genome was mapped to a similar depth and resolution, it would predict that most, if not all, promoters and enhancers would form multi-way interactions. However, in the *Ppm1g* and *Klf1* loci, multi-way interacting promoters are not co-regulated by the same co-activators (**Supplemental Figure 9**). Similarly, multi-promoter interacting enhancers identified by RIC-seq show stark differences in CBP/p300-dependency.

What could be the reasons for extensive multi-way E-P interactions but a lack of coregulation of the interacting promoters? This could be due to the property of accessible chromatin regions to interact with each other or enhancers interacting with many promoters but activating only specific ones based on biochemical compatibility ^52,62^. Regardless of the reason, our results suggest two limitations in inferring functional relationships from physical E-P contacts: (1) extensive multi-way E-P interactions identified in deep, locus-specific analyses appear to be poor indicators of functional gene regulatory relationships, and (2) genome-scale E-P interaction maps underestimates the role of short-range enhancer interactions, which are highly consequential for gene regulation according to CRISPRi data.

Despite these limitations, we emphasize that studying chromatin interactions remains valuable for understanding the principles of genome folding and elucidating the mechanistic details of looping between functionally characterized E-P pairs. However, using E-P interaction mapping as a discovery tool for genome-scale identification of enhancer and GWAS targets, as done routinely in the current literature, needs careful reevaluation. At a minimum, gene-skipping and non-skipping enhancer targets identified in genome-scale E-P contact mapping analyses should be considered as distinct groups, and active gene-skipping enhancer targets should be treated with lower confidence.

To be clear, we are not disputing the well-established model in which enhancers physically contact promoters and activate genes over very long distances (>1 Mb) ^1–3,7,11–14^. Rather, we concur with this model and our own concurrent work shows that cohesin-dependent looping promotes the activation of hundreds of genes in mammalian cells ^63^. What this work challenges is the idea that a large portion of mammalian enhancers intentionally skips over active genes to activate distant targets, or that a single enhancer often activates multiple promoters in a coordinated group, as suggested by multi-way interactions in RCMC and RIC-seq ^28,34^. Instead, we propose that enhancers do not typically use chromatin looping to deliberately bypass neighboring active genes; instead, looping mostly serves as an efficient mechanism to reach out to neighboring genes that reside at longer genomic distances, and in rare instances, to skip active genes.

### CRISPRi and eQTL analyses likely systematically underestimate distal enhancer contribution

Each method has its strengths and limitations. While considering the limitations of 3C-based analyses, it’s also important to mention the limitations of CRISPRi. Our findings suggest that CRISPRi has an opposite bias to Micro-C and HiC, which results in a likely overestimation of the contribution of proximal enhancers and an underestimation of the contribution of distal enhancers. If enhancers reside very close (<3kb) to promoters, CRISPRi-induced repression may extend to promoters and result in overestimating enhancer contribution.

The underestimation of distal enhancer contribution likely occurs because gene expression measurement methods are inherently biased for detecting strong effects. On a genome-wide scale, the gene activation ability of native CBP/p300-dependent enhancers decreases with distance ^50^, which is consistent with synthetic p300 targeting and enhancer insertion studies ^64,65^. As a result, CRISPRi more easily detects proximal enhancers that cause strong gene regulation changes, while distal enhancers, which have weaker effects, are often missed.

This bias likely extends to the discovery of eQTLs. The discovery of GWASs and eQTL is affected by distinct parameters. GWAS variant discovery is influenced by their frequency in the population and the strength of their association with traits, rather than their distance from gene promoters. As a result, both promoter-proximal and distal GWAS variants can be identified equally well. However, eQTL discovery relies on detecting changes in mRNA expression, making it more biased toward identifying targets of proximal GWAS variants causing strong expression effects. This bias may result in missing the targets of distal variants, despite their biological relevance. This may, at least partially, explain the reduced overlap between distal GWAS variants and eQTLs. While factors like trans-eQTLs, sample size, and cell-type specificity contribute to this ^66–69^, they alone dońt fully explain the lower overlap of distal GWASs as compared to proximal ones. Our results showing that enhancers rarely skip active genes and their function decreases with distance can help explain why most (∼95%) lead cis-eQTL SNPs are found within 100 kb of their target genes ^70^.

### CBP/p300-dependent enhancers are the predominant type, and they seldom skip active genes

Mammalian genes are regulated by different types of enhancers ^52,58–60,71^. Defining the nature of different enhancer types and identifying the predominant enhancer type is a prerequisite for understanding the intricacies of gene regulation. Our analysis of approximately 800 CRISPRi-identified enhancers and GWAS-gene pairs has identified H2BNTac-positive, CBP/p300-dependent enhancer as the predominant enhancer type in mammalian cells. Enhancers of this type have the following attributes: they are marked by H2BNTac, drive robust gene activation, preferentially activate cell-type-specific genes, and their regulatory influence decreases with distance. Enhancers of this type mainly activate neighboring genes, and ∼5% of them may impact multiple proximal genes, mostly located consecutively. This enhancer type can skip active genes, but this is an exception rather than a rule. For this enhancer type, we estimate that >90% have targets within the first neighboring active genes, >95% within two neighboring genes, and ∼99% within three neighboring genes (**Figure 4F**). For this enhancer type, we believe that our approach, based on linear E-P proximity and CBP/p300 dependency of proximal genes, can predict enhancer targets with accuracy comparable to, or better than, CRISPRi. Because of its accuracy, it can effectively distinguish between gene-skipping and non-skipping enhancers in CRISPRi datasets.

### CRISPRi-identified enhancers functioning without CBP/p300 have atypical properties

What are the attributes of other enhancer types, and what is their regulatory scope? Among genome-scale CRISPRi-defined E-G pairs, around a quarter of enhancer target genes are activated without CBP/p300 and they may represent other (non-CBP/p300-dependent) enhancer types. These enhancers have the following traits: weak or absent H2BNTac, no preference for cell-type-specific genes, frequent skipping of active genes, and weak gene downregulation with poor FDR in CRISPRi assays. Strikingly, the proportion of these enhancers increases linearly with the broadening of the enhancer target search window, suggesting they operate without any distance constraints. Do these enhancers rely more heavily on cohesin-dependent looping when contacting their distal targets? The answer seems to be NO. Our concurrent work shows that acute cohesin depletion mostly downregulate genes that are activated by CBP/p300 ^63^.

Given these atypical attributes, two factors must be considered in interpreting their role in gene regulation: (1) Approximately 10% of E-G pairs in the CRISPRi dataset are likely false positives based on the FDR. (2) Gene expression changes were measured 10 days post-CRISPRi introduction, suggesting that some identified targets may be secondary or indirect. Our analyses show that, at a fixed p-value, false positives increase linearly with the broadening of target search windows. At a controlled FDR, the number of identified significant enhancer targets decreases. For example, at an FDR of 1%, the number of significant enhancer targets drops fourfold when the window is expanded from 100 kb to 500 kb. At an FDR of 5%, significant targets decrease by over 50% when extending the search from 200 kb to 1 Mb. Within FDR-controlled data, it is difficult to discern true positives from false positives and indirect enhancer targets. However, false positives and indirect targets would predictably lack cell-type specificity, show no CBP/p300 regulation bias, and become more prevalent as the target window broadens. We do not rule out the existence of long-range acting enhancers functioning without CBP/p300. However, because the CRISPRi-identified CBP/p300-independent enhancers exhibit unusual traits, their authenticity should be interpreted cautiously.

In summary, we uncover clear and systematic differences between gene-skipping and non-skipping enhancers identified by genome-scale approaches. Regardless of the underlying reasons, our findings have major implications for evaluating the use of different approaches for identifying enhancer targets, and for predicting enhancer targets based on distance and coactivator dependencies. While our conclusions are based on a single cell type, the conserved nature of gene regulation principles would suggest that these principles likely apply to other mammalian cell types. Even if they do not, it does not diminish the fact that enhancer targets identified by different methods systematically differ in the widely used K562 model cell line.

## Supplemental Notes

### Supplemental Note 1

To infer if CBP/p300-dependent enhancers truly skip proximal active genes, we manually assessed the genome browser tracks of 18 ^highconf^E-G pairs where the enhancers skipped one or more proximal active genes, and the enhancer-linked distal target genes were downregulated (HD+ID) by A-485 (**Supplemental Table S1B)**. The A-485-induced downregulation of their distal target genes suggests these enhancers function using CBP/p300. If these enhancers indeed skipped proximal genes, then we would anticipate that their nearest genes will remain unaffected by A-485. Furthermore, if the CRISPRi-nominated enhancers regulate the distal targets, then those distal target genes will not be anticipated to harbor other H2BK20ac-marked enhancers in their proximity.

Of the 18 enhancers, in our manual interpretation, 2 enhancers (linked to *TPST2* and *ALAS2*) do not skip active genes because the skipped genes are not detectably expressed in nascent transcriptome analyses, and 3 other enhancers (linked to *AKR1C2*, *DECR1*, and *ELK1*) activate consecutively located genes, without skipping an intervening active gene. For example, the enhancer linked to *TPST2* likely does regulate this gene, but it does not appear to skip an active gene because the two intervening genes, *CRYBA4* and *CRYBB1*, are not detectably expressed. The enhancer linked to *DECR1* skips *CALB1*, but the CBP/p300 dependency of these genes indicates that this enhancer likely mainly regulates the proximal genes, *LINC00534* and *CALB1*, and even if it impacts the distal target, *DECR1*, the effect is likely weak.

For the remaining 13/13 enhancers, each enhancer had at least one proximal gene that was downregulated by A-485, but for 9 of these enhancers, the A-485 downregulated enhancer proximal genes did not score significantly in CRISPRi analyses. Yet more notably, 13/13 distal targets of these enhancers had strong H2BK20ac marked proximal enhancers, indicating that the A-485 sensitivity of the distal genes probably reflects the impact of the proximal enhancers that are distinct from CRISPRi-identified enhancers. For example, the enhancer linked to *CYSTM1* is more likely to regulate the proximal gene *CXXC5* instead, and *CYSTM1* is likely regulated by other enhancers that occur proximal to this gene.

Overall, manual inspection confirms that several of the gene-skipping 18 ^highconf^E-G pairs linked to A-485-downregulated genes likely regulate multiple consecutively located proximal genes, but we did not find unambiguous evidence of any of these enhancers entirely skipping proximal active genes to regulated distal genes exclusively.

### Supplemental Note 2

We manually examined the genome browser tracks of the significant scoring 664 E-G pairs in the Gasperini et al. dataset for the presence of H2BK20ac in targeted enhancers, presence of non-CRISPRi targeted H2BK20ac-positive regions, E-G proximity, A-485-induced regulation of the enhancer proximal genes in K562, and in some instances in additional human cell lines (two blood cell lines: Jurkat, KG1; and 3 non-blood cell lines: HepG2, HCT116, SH-SY5Y). Based on our manual assessment, we grouped E-G pairs into three groups. E-G pairs in Group 1, Group 2, and Group 3 have the following characteristics.

Group 1 includes 64.3% (427/664) E-G pairs from the entire dataset. 78.4% of E-G pairs in Group 1 are from the high-confidence group, and 21.6% are from the low-confidence group. In Group 1, 99% of enhancers are marked by H2BK20ac, 97% rank in the top 10,000 H2BK20ac enrichment, and 1.5% of E-G pairs have a distance >200kb. From the unbiased computer annotations, 4% (17/428) of Group 1 E-G pairs involve gene-skipping. Manual inspection of these 17 E-G pairs shows that 8 of them include enhancers that activate consecutively located, proximally occurring genes without avoiding activation of any active promoter, and 6 E-G pairs skip genes that are not detectably expressed in K562. The remaining 3 E-G pairs (two linked to *MPST* and one linked to *IFITM1*) may genuinely skip active promoter(s) without activating them.

Group 2 includes 26.7% (177/664) E-G pairs from the entire dataset. In this group, 56.5% of E-G pairs are from the high-confidence group, and 43.5% are from the low-confidence group. In Group 2, 43% of enhancers have H2BK20ac rank >10,000 or lack detectable H2BK20ac, and 85.5% of E-G pairs involve gene skipping. Just 4.5% of E-G pairs in this group have enhancers that rank in the top 10,000 and are liked to proximal genes (non-skipping). In Group 2, 12.8% of E-G pairs involve A-485 downregulated (HD+ID) genes. However, all of the A-485 downregulated genes in this group have strong H2BK20ac marked proximal enhancers that are distinct from the enhancers identified by CRISPRi, and the occurrence of proximal H2BK20ac-positive enhancers can best rationalize the A-485-induced downregulation of these genes. Thus, Group 2 enhancers are characterized by a high frequency of skipping proximal genes, weak or no H2BK20ac, and lack of requirement for CBP/p300 for their target gene activation.

Group 3 includes 9% (60/664) of E-G pairs, which we could not categorize into Groups 1 and 2. In Group 3, 58% of pairs are from the high-confidence group, and 42% are from the low-confidence group. In this group, CRISPRi-identified enhancer targets were either not quantifiable due to low expression (36.7%), or CRISPRi-identified enhancers were marked by H2BK20ac, but A-485 did not regulate their targets, and the reason for A-485 insensitivity could not be rationalized from analyzed chromatin features.

Overall, we conclude that H2BK20ac-positive, CBP/p300-dependent enhancers can regulate multiple genes, but they rarely (∼1-2%) skip proximal active genes to regulate distal genes exclusively. We recognize that manual annotations can be prone to bias; therefore, all annotations are provided in the supplemental data (**Supplemental Table S1c**) so that readers can make independent assessments.

### Supplemental Note 3

To test whether H2BK20ac enrichment could predict enhancer strength, we compared CRISPRi-determined GWAS strength with H2BK20ac-predicted strength. In 47 genes, CRISPRi targeted multiple proximal GWASs, with some scoring significant and others not (**Supplemental Table S2b**). For each of these genes, up to 3 strongest GWASs were ranked by their strength, determined empirically by CRISPRi-induced downregulation of their nearest genes. The same GWASs were ranked by their predicted strength based on H2BK20ac enrichment. Then, the concordance between CRISPRi-determined and H2BK20ac-predicted GWAS ranking was checked. This analysis included 115 GWASs, encompassing more than half (69/128) of the significant scoring GWASs in the Morris et al. dataset. For two-thirds of genes (33/47 genes; with 83 proximal GWAS), H2BK20ac enrichment correctly predicted the CRISPRi-defined strongest GWASs (concordant match), but for one-third of genes (14/47 genes, with 32 proximal GWASs), H2BK20ac enrichment failed to correctly predict the CRISRI-defined strongest GWAS (discordant match) (**Supplemental Figure 5A-B**).

The concordant (n= 33) and discordant (n= 14) matching GWASs show notable differences (**Supplemental Figure 5C**). (1) Discordant matching GWASs have weak H2BK20ac; 8/14 discordant GWASs had ^low,no^H2BK20ac, whereas only 3/32 concordant matching GWASs did. (2) Discordant matching GWASs caused very weak gene downregulation in CRISPRi. 9/14 discordant GWASs had CRISPRi log2FC > - 0.4 (remaining transcripts >75.8%), whereas only 5/32 concordant GWASs caused such weak downregulation. (3) Discordant GWASs are biased for promoter proximity. Four of the five strongest discordant GWASs occurred within 3kb of TTSs, whereas among the five strongest concordant GWASs, only one occurred within this distance.

These analyses demonstrate that in most cases, H2BK20ac enrichment correctly ranks the relative function of gene proximal GWASs. However, exceptions include CRISPRi-identified GWASs causing very weak gene downregulation and those located very close to the TSS of their target genes. These exceptions possibly result from inaccuracies in quantifying weakly downregulated targets and overestimating GWAS impact by CRISPRi when targeted GWASs are close to gene promoters.

### Supplemental Note 4

To understand why some H3BK20ac-positive GWASs score significantly while others dońt, we focused on 247 GWASs that ranked in the top 10,000 H2BK20ac peaks and are presumably active. Of these, 80 GWASs scored significant (^sig^GWASs), and 167 scored not significant (^nonsig^GWASs) in the CRISPRi analysis.

Significant and non-significant scoring GWASs differed in several ways. In comparison to ^sig^GWASs, ^nonsig^GWASs display weaker chromatin accessibility (p= 1.1e-6, Mann-Whitney U-test) (**Supplemental Figure 6A**), exhibit weaker H2BK20ac enrichment (p= 0.02, Kolmogorov-Smirnov test) (**Supplemental Figure 6B**), GWAS targeting gRNA bind further away from the nearest ATAC-seq peak center (**Supplemental Figure 6C**), and notably, they are located farther from their nearest active genes (median distance: ^sig^GWAS, 11.2kb; ^nonsig^GWAS, 26.8kb, p= 7.2e-4, Mann-Whitney U-test) (**Supplemental Figure 6D**). Moreover, within ^nonsig^GWASs, those occurring proximally to SD+ND genes exhibit greater distances than those proximal to HD+ID genes (median distance 18.6kb versus 40.5kb, p= 5.6e-4, Mann-Whitney U-test).

Despite these differences, multiple lines of evidence suggest that many ^nonsig^GWASs still exert a functional impact on their nearest genes. While ^nonsig^GWAS proximal genes are individually not significantly regulated by CRISPRi, overall, they show a statistically significant tendency of downregulation (^nonsig^GWAS proximal versus non-proximal gene, p= 7.8e-16, Mann-Whitney U-test) (**Supplemental Figure 6E**). ^nonsig^GWAS proximal genes include a much higher fraction of HD genes than all genes expressed in K562 (41% versus 9.5%) (**Supplemental Figure 6F**). The extent of CRISPRi-induced downregulation of ^nonsig^GWAS proximal genes is associated with the extent of their A-485-induced downregulation (**Supplemental Figure 6G**). CRISPRi-induced downregulation of ^nonsig^GWAS proximal genes is related to the enrichment of H2BK20ac level in ^nonsig^GWASs (**Supplemental Figure 6H**). These results suggest that many ^nonsig^GWASs contribute to regulating their proximal genes. However, their quantitative impact is subtle compared to ^sig^GWASs, and CRISPRi analysis is underpowered to detect these small changes. We postulate that the same limitation also hampers identifying eQTLs for weak GWASs, prominently including distal GWASs.

## Acknowledgments

We thank the members of the Choudhary lab for their helpful discussions. The Novo Nordisk Foundation Center for Protein Research is financially supported by the Novo Nordisk Foundation (NNF14CC0001). C.C. is supported by the Novo Nordisk Foundation Distinguished Investigator Bioscience and Basic Biomedicine (NNF22OC0074677). We thank Dr. Yuanchao Xue for sharing RIC-seq identified enhancer-promoter interaction dataset.

## Author contributions

Project conceptualization and funding acquisition, C.C.; research design, C.C. and T.N.; data analysis and figure preparation, T.N.; manuscript writing C.C. and T.N.

## Code availability

All software and packages used for data analysis are described wherever relevant in the Methods and Reporting summary.

## Methods

### Analysis of EU-seq nascent transcription data

The EU RNA-seq data (GSE146328 and GSE196250) treated with either DMSO or A-485 (10 µM, treatment time of 60 min in human cell line, and 30 min, 60 min, and 120 min in mouse ESC) were reprocessed. Adapters and low-quality sequences (Phred quality score < 20) were removed using Cutadapt v.4.2 (https://doi.org/10.14806/ej.17.1.200). The read sequences were aligned to the human hg19 (GENCODE Grch37 release 29) or mm10 (GRCm38 version 6) utilizing the bwa aln tool with default parameters (BWA v.0.7.10)^72^. Reads with multiple mappings and those with more than three mismatches were filtered out using samtools (v1.4)^73^. The ribosomal and transfer RNA regions were retrieved from the UCSC genome browser and reads aligning to these regions were excluded using BEDTools (v.2.23)^74^. The nascent transcription changes were calculated by using genebody regions of maximum 30kb, 90kb and 240kb for the A-485 treatment data at 30 min, 60min and 120 min, respectively, based on the polymerase II elongation rates. The reads mapped to the defined gene body regions were counted using HTSeq (v.0.11.1)^75^. Only the transcript types ”protein-coding” and ”lincRNA” were considered for building the active gene model. The isoform selection across each cell line consisted of two steps. In the first step, highly confidently expressed transcripts were chosen based on three criteria: 1) transcripts per million (TPM) > 2 in RNA-seq, 2) the identification of H3K4me3 ChIP-seq peak within 1kb of the transcription start site (TSS), and 3) an average EU-seq read count per replication and condition exceeding 20. When multiple isoforms remained, the longest isoform was selected. In the second step, for protein-coding or lincRNA genes not included in the first step, the longest transcripts were included if they had an average mapped read count greater than 20 per replicate and condition in EU-seq analyses. Log2 fold-changes (FC) after A-485 treatment were calculated using DEseq2(v.1.32.0)^76^. In mouse ESC, Log2 fold-changes (FC) after A-485 treatment were calculated by averaging log2 fold-changes measured at three time points. In the calculation of scaling factors, we used read counts only from highly confidently expressed transcripts (i.e., transcripts selected in step 1) and calculated by the default DEseq2 method, which normalizes the read counts using the median of their relative abundance across conditions and replicates. The regulation by A-485 was categorized based on the remaining transcripts as follows: ND: > 83.3% (<1.2-fold downregulation), SD: 66.7 - 83.3% (1.2-1.5-fold downregulation), ID: 50 - 66.7% (1.5-2-fold downregulation), HD: ≤ 50% (≥2-fold downregulation).

### Processing of RNA-seq data

Mouse and human cell line RNA-seq data were re-processed from pre-deposited data (GSE196250). Adaptors and low-quality sequences (-q 20) were trimmed using Cutadapt (https://doi.org/10.14806/ej.17.1.200). Reads were aligned to the human hg19 (GENCODE Grch37 release 29) or mm10 (GRCm38 version 6) using STAR (v2.6.1a)^77^ with the removal of the non-canonical junctions. Reads mapped to protein-coding or lincRNA exon were counted on a gene basis using HTseq (v.0.11.1)^75^. TPM was calculated using R.

### Processing of ChIP-seq data

Reads were aligned to the mm10 and human hg19 genome by using bwa mem with default parameters (BWA version 0.7.10). Multi-mapped, duplicated, or reads with more than three mismatches were filtered out by samtools^73^. Reads mapped to the DAC Blacklisted Regions (https://www.encodeproject.org/annotations/ENCSR636HFF/) were excluded from further analysis. Peak regions were identified using LanceOtron with the default model (wide-and-deep_jan-2021)^78^. Peak height was determined using bamCompare^79^ with the following parameters (centerReads, minMappingQuality 10, 20bp bin, smooth length 400bp, extend reads 200bp, rpm normalization, and subtraction of the input rpm value). The peaks proximal within a proximity of 2kb were merged using Bedtools^74^, and poorly enriched peaks of peak summit height < 8 reads mapped per million (rpm) were discarded.

### H2BK20ac enrichment analysis of gRNA target regions

Initially, we generated a rank list for H2BK20ac enrichment in candidate enhancer regions. In each called H2BK20ac peak region, we counted H2BK20ac reads within a +/- 1Kb window around the peak summit, normalized to rpm, and input rpm was subtracted. Where specified, reads mapping near TSS (+/- 500b regions) were excluded to avoid the potential contribution of promoter acetylation. Peaks with ≥1 rpm were retained to define H2BK20ac marked candidate enhancers. Using these parameters, we identified a total of 20,408 (K562), 22,164 (KG-1), and 19,029 (Jurkat) candidate enhancer regions. The identified candidate enhancers were ranked according to their H2BK20ac enrichment in the respective cell lines, used as a reference for determining H2BK20ac enrichment-based ranking of CRISPRi targeted enhancers and GWASs and RIC-seq identified enhancers.

Next, H2BK20ac enrichment of gRNA target regions (for CRISPRi data) and enhancer regions (for RIC-seq data) were calculated using the following regions: Gasperini et al. the center of gRNA target region +/- 1Kb, Morris et al. GWAS position +/- 1Kb, and Liang et al. the highest enhancer peak summit +/- 1Kb. The H2BK20ac enrichment of these regions was computed using the same procedure applied to the H2BK20ac peak regions. If the gRNA target region or RIC-seq enhancer region overlapped with the identified H2BK20ac candidate enhancer peaks, the H2BK20ac enrichment rank for the gRNA target regions and RIC-seq enhancer regions was determined using H2BK20ac peak ranks. If the gRNA target region or RIC-seq enhancer region has an H2BK20ac rpm value that is lower than the H2BK20ac rpm value of *jth-*ranked reference enhancer but greater than that of (*j + 1)* th ranked reference enhancer, then it is ranked as *j+1*. If the gRNA target region or enhancer region did not overlap with the identified H2BK20ac peaks, it was classified as “None.” Consequently, enhancers in K562 were ranked from 1 to 20,409, or “None.”

### Processing of ATAC-seq data

K562 and HCT116 ATAC-seq raw data was downloaded from the NCBI GEO database (GSE97889, and GSE213909) and reprocessed. Paired reads were aligned to the hg19 genome using BWA meme (version 1.0.4)^80^. Paired reads were aligned to the hg19 genome using BWA meme (version 1.0.4)^80^ with the soft clipping option for supplementary alignments. Duplicated read pairs were removed using Picard-tools (version 2.9.1, “Picard Toolkit.” 2019. Broad Institute, GitHub Repository. https://broadinstitute.github.io/picard/; Broad Institute). We filtered out low-quality reads with a MAPQ score of less than 10 and non-primary alignments and retained only the properly aligned read pairs using samtools. Peak regions were identified using LanceOtron with the default model (wide-and-deep_jan-2021)^78^. Peak height was calculated using bamCoverage with the following parameters (centerReads, 20bp bin, smooth length 400bp, extend reads 200bp, rpm normalization). Poorly-enriched peaks of peak summit height < 8 rpm were filtered out. For the calculation of distance between SNPs and ATAC-seq peak summit, we considered only ATAC-seq peaks that overlapped with H2BK20ac peak regions.

### Cell type specificity of CRISPRi significant genes

We used the FANTOM5 CAGE dataset as a reference to determine the degree of cell-type specificity of certain genes^46,47^. A binary expression matrix was constructed from 76 human tissue expression profiles (comprising 68 tissues from adult and fetal stages), applying a gene expression threshold of TPM ≥ 2. We then plotted a cumulative fraction of K562 expressed genes and CRISPRi-defined enhancer target genes in relation to the number of tissues expressing these genes.

### Analysis of CRISPRi data from Gasperini et al

The processed CRISPRi data on K562 were retrieved from the GEO repository under accession GSE120861 as a supplementary file (GSE120861_all_deg_results.at_scale.txt.gz). We adhered to the active gene model and transcription start sites (TSSs) as determined by Gasperini et al.^40^. Outlier genes, defined by Gasperini et al., which exhibited higher expression levels than anticipated and presented difficulties in accurate expression changes calculation were excluded from the analysis. We employed the "top_two" groups of gRNAs and designated "DHS" as site types in the datasets. Consequently, 5,729 candidate cCREs and 78,440 E-G pairs were retained for the subsequent analysis. The distance from TSS to the center of gRNA target sites was used to determine enhancer-promoter distances. The 470 high-confidence and 194 low-confidence E-G pairs defined by Gasperini et al. were utilized as significant E-G pairs. Additionally, 54,071 E-P pairs exhibiting CRISPRi log_2_FC > -log_2_(1.05) were categorized as non-significant. To analyze H2BK20ac enrichment, we adopted the center +/- 1Kb region of gRNA target sites and assessed their enhancer ranks as described in the “H2BK20ac Enrichment Analysis of Enhancer and SNP Regions” section.

To examine the impact of varying target search windows, we first calculated the nominal p-values for E-G pairs encompassing a 20Mb distance window and non-targeting controls (NTCs) connected with all the active gene pairs. We utilized processed scRNA-seq data and get_deg.at_scale.R function available on the github repository (https://github.com/shendurelab/tafka-crisprQTL). The reference list of p-values was generated from NTC-all active gene pairs. The nominal p-values of E-G pairs are converted to empirical p-values using the reference p-value list using the following formula described in Gasperini et al. Empirical p value = ((the number of NTCs with a smaller nominal p value than that test’s nominal P value) + 1) / (the total number of NTCs tests + 1). In the analysis examining the impacts of various target search windows, we adjusted the empirical p-values by considering all the p-values present within these windows. E-G pairs were categorized as significant under each FDR threshold, with corrected empirical p-value falling below the designated threshold of 0.1, 0.05, or 0.001. In the analysis using the fixed p-value threshold, we implemented an empirical p-value threshold of 2.77e-3, corresponding to an FDR of 0.1 within 1Mb target search windows. Across different target search windows, E-P pairs falling below this threshold were considered for analyses.

### Analysis of STING-seq data from Morris et al

The processed data from Morris et al. were downloaded as a supplementary table (Table S3F). The GWAS-gene pairs with ≤ 1Kb were excluded. As a result, 7,981 GWAS-gene pairs with 539 GWAS were utilized for further analysis. Unless otherwise mentioned, 148 down-regulated GWAS-gene pairs with Q values < 0.05 within a 500Kb target search window were classified as significant, and the remaining pairs were categorized as non-significant. For the calculation of H2BK20ac enrichment, GWAS +/- 1Kb regions were used and ranked as described in the “H2BK20ac and H3K27ac enrichment analysis of gRNA target regions” section.

### Analysis of RIC-seq data from Liang et al

The dataset of RIC-seq identified enhancers and active promoter group regions^28^ was kindly shared by Dr. Yuanchao Xue through personal communication. The regions of enhancer and promoter groups identified in the RIC-seq dataset were broader than those in the CRISPRi datasets. Thus, we determined the E-P distances based on an end-to-end basis. E-P pairs located on different chromosomes were categorized as “inter-chromosomal.” To calculate the enrichment of H2BK20ac in enhancers, we used the highest peak summit region +/- 1Kb within each enhancer. The procedure for defining H2BK20ac rank is described in the “H2BK20ac enrichment analysis of gRNA target regions” section. Furthermore, we explored ranking enhancers using all the enhancer regions for H2BK20ac calculations and found negligible differences in overall analysis results. Of the 3,780 unique promoter groups in the RIC-seq datasets, 1,944 promoter groups (51.4%) comprise multiple genes. Unless indicated otherwise, we selected the gene most down-regulated by A-485 as its representative for each of these promoter groups.

To investigate if enhancers linked to multiple promoters activate genes using the same coactivators, we selected RIC-seq-identified enhancers that interact with multiple promoter groups, with at least one promoter corresponding to a gene downregulated by more than 2-fold after A-485 treatment. After excluding the most regulated genes, based on which the multi-gene-linked enhancers were selected, we then examined the A-485-induced regulation of other genes.

### Processing of GWAS data from Morris et al

Fine-mapped blood cell trait GWAS variants from the UK Biobank (UKBB) and Blood Cell Consortium (BCX) were downloaded from Morris et al. (Table S1, C and D)^41^. In analyzed cell lines, H2BK20ac enrichments were determined in GWAS +/- 1Kb regions, and subsequently, GWASs were ranked as described in the “H2BK20ac and H3K27ac enrichment analysis of gRNA target regions” section. The active genes and TSS positions in each cell line were determined by using H3K4me3 peak positions, RNA-seq data and EU-seq data as described in the “Nascent transcription analyses by EU-seq” section.

### Manual annotation of CRISPRi identified E-G pairs

We manually examined the genome browser tracks of the significant scoring 664 E-G pairs in the Gasperini et al. dataset. We checked for the presence of H2BK20ac in targeted enhancers, the presence of non-CRISPRi targeted H2BK20ac-positive regions, E-G proximity, A-485-induced regulation of CRISPRi identified target genes, and enhancer proximal genes in K562, as well as in 5 other human cell lines (Jurkat, KG1, HepG2, HCT116, SH-SY5Y).

Each E-G pair was evaluated individually, and observations from manual assessments are provided in Supplemental Table S1C. Based on the observed properties, the pairs were grouped into three categories:

Group 1: Likely or plausible targets of CRISPRi-identified enhancers. Likely enhancer targets are defined as E-G pairs that include enhancers marked by H2BK20ac-positive in manual inspection and show indication of regulation by CBP/p300. Plausible enhancer targets include pairs with H2BK20ac-positive enhancers and some indication of CBP/p300 regulation in K562 or other human cell lines.

Group 2: E-G pairs where CRISPRi-identified enhancers have weak or no H2BK20ac enrichment, and/or the target genes are not regulated by CBP/p300. This group also includes pairs involving active gene skipping, where the CRISPRi-identified distal target gene is regulated by CBP/p300 but better explained by proximal enhancers other than those identified by CRISPRi.

Group 3: E-G pairs where the CRISPRi-identified enhancer targets that are expressed at a very low level in nascent transcription data, and the regulation of genes could not be confidently evaluated in manual inspection. This group also includes pairs where CRISPRi-targeted enhancers are marked by strong H2BK20ac, but the identified enhancer targets are not regulated by CBP/p300, and we could not rationalize why these genes are not affected by CBP/p300 inhibition.

### Annotation of SE-regulated genes

The MED1-enriched super-enhancer regions in mouse ESC were downloaded from dbSUPER ^81^, and converted to mm10 by using UCSC liftover tools ^82^. To identify SE-regulated candidate genes, we manually examined genome browser tracks to identify the nearest active genes based on nascent transcript expression in previously published EU-seq data in mESC^83^. In some instances, we found that the nearest active genes encode transcripts of currently unannotated genes. If the expression of SE proximal active gene(s), including unannotated genes, was downregulated in A-485 treated cells, the genes were considered candidate SE target genes. A-485-induced fold-change in all candidate SE target genes was determined using nascent transcript inhibition by A-485. The regions used for calculating A-485-induced transcription change in unannotated transcripts are specified in Supplemental Table S5. If neither of the SE neighboring genes were downregulated by A-485, we deemed that SE does not regulate proximal gene. In these instances, we calculated A-485-induced fold-change for the gene nearest to SE.

### Processing of MPRA data in HCT116

The processed differential analysis data of COF-AID STARR-seq was downloaded from Neumayr et al. ^52^. COF-dependent enhancer regions were identified based on either by 1) decreased enhancer activity following COF depletion with an FDR < 0.05, or 2) > 1.5-fold decrease in enhancer activity and were analyzed separately. COF-independent (N.C) enhancers were defined by having absolute changes after COF depletion < 1.2-fold and an FDR > 0.5. COF-dependent enhancers were further categorized from Q4 to Q1 based on the degree of regulation from highest to lowest within each experiment. STARR-seq enhancer regions were characterized by their overlap with ATAC-seq or H2BK20ac peaks, along with their intensities. The ATAC peak height within each enhancer region was determined using the highest ATAC-seq rpm value calculated as described in the Processing of ATAC-seq data section. H2BK20ac enrichment was determined using the input subtracted H2BK20ac rpm values in the corresponding enhancer regions.

### Processing of Micro-C data

K562 Micro-C raw data were downloaded from GSE206131 and reprocessed ^42^. Reads were mapped to hg19 genome by using the distiller-nf v0.3.4 pipeline (https://github.com/open2c/distiller-nf). The paired-end reads were aligned to the hg19 genome by using BWA mem ^72^. Pairs with low mapping quality (mapq of read1 and/or reads2 < 30) were filtered out. PCR duplicates were identified by allowing maximum of three mismatches in the mapping regions and were subsequently filtered out. Contact regions were identified on the KR-balanced Micro-C data using hiccups (juicer tools, version 1.22.01) ^45^ at 1kb, 2kb, and 5kb bin resolutions. The following parameters were used for calling enriched pixels: window width; 10kb (1kb resolution), 16kb (2kb resolution), 30kb (5kb resolution), peak width; 4kb (1kb resolution), 8kb (2kb resolution), 15kb (5kb resolution), distance used for merging proximal pixels; 2.5kb (1kb resolution), 5kb (2kb resolution), 10kb (5kb resolution). Enriched contacts with an FDR < 0.1 from the three resolutions were merged, ensuring that if contacts from lower-resolution data overlapped with those from the higher-resolution data, only contacts from the higher resolution were retained. As a result, a total of 28,697 merged Micro-C loops were used for annotating CRISPRi data. CRISPRi E-P pairs from Gasperini et al. were considered to overlap with Micro-C loops if they matched within a 1kb distance.

### Annotation of RCMC data

The RCMC enhancer-promoter and promoter-promoter loop data were kindly provided by Dr. Anders S Hansen. The loop regions were converted from mm39 to mm10 by using UCSC liftover tools ^82^. Promoters of actively transcribed genes were identified based on the presence of the promoter mark H4K4me3 and nascent transcript expression patterns. Candidate active enhancers were identified using H2BK20ac marked ATAC-seq peaks, and potential enhancer targets were inferred based on the genes’ reliance on CBP/p300. In our manual annotations, a few of the elements that were originally classified as candidate “enhancer” were re-classified as “promoters,” and vice-versa, a few of the elements that were originally classified as candidate “promoters” were re-classified as “enhancers,” as indicated in Supplemental Figure 9. The re-classification was based on the presence of H3K4me3, H2BNTac, and nascent transcript expression patterns. The E-P and P-P contacts shown in Supplemental Figure 9 are based on the original manual annotations shared by Goel et al ^34^. Enhancer candidate target genes in the RCMC analyzed loci were identified based on A-485-induced nascent transcription changes as described above for identifying SE candidate target genes.

### Statistical analysis

Unless otherwise mentioned, p-values are calculated using the two-sided Mann–Whitney U test. Multiple comparisons were corrected using the Benjamini & Hochberg method (R package stats version 3.6.2).

### Use of AI tools

The manuscript was written by authors. ChatGPT 3.5 was used for checking grammatical errors.

### Use of publicly available data

Human H2BK20ac and human EU-seq are generated as described in references^48,50^. The following datasets were also downloaded and analyzed: Human and mouse reference genome and annotation from the GENCODE website (human; Grch37, version 29, mouse; GRCm38 version 6). ENCODE DAC Blacklisted Regions (https://www.encodeproject.org/annotations/ENCSR636HFF/); K562 CRISPRi data from Gasperini et al.^40^ (supplementary data GSE120861_all_deg_results.at_scale.txt, and processed scRNA-seq data and gRNA data from https://github.com/shendurelab/tafka-crisprQTL); K562 STING-seq data (Table S3, F), UKBB and BCX blood cell traits GWAS data from Morris et al.^41^ (Supplementary Tables S1, C and D), COF-AID STARR-seq from Neumayr et al ^52^ (Supplementary Table 3). RIC-seq data and active promoter group position were kindly shared by Dr. Yuanchao Xue through personal communication. RCMC contact data and their contact type annotation were kindly shared by Dr. Anders S Hansen through personal communication.

---

**Supplemental Figure 1.**
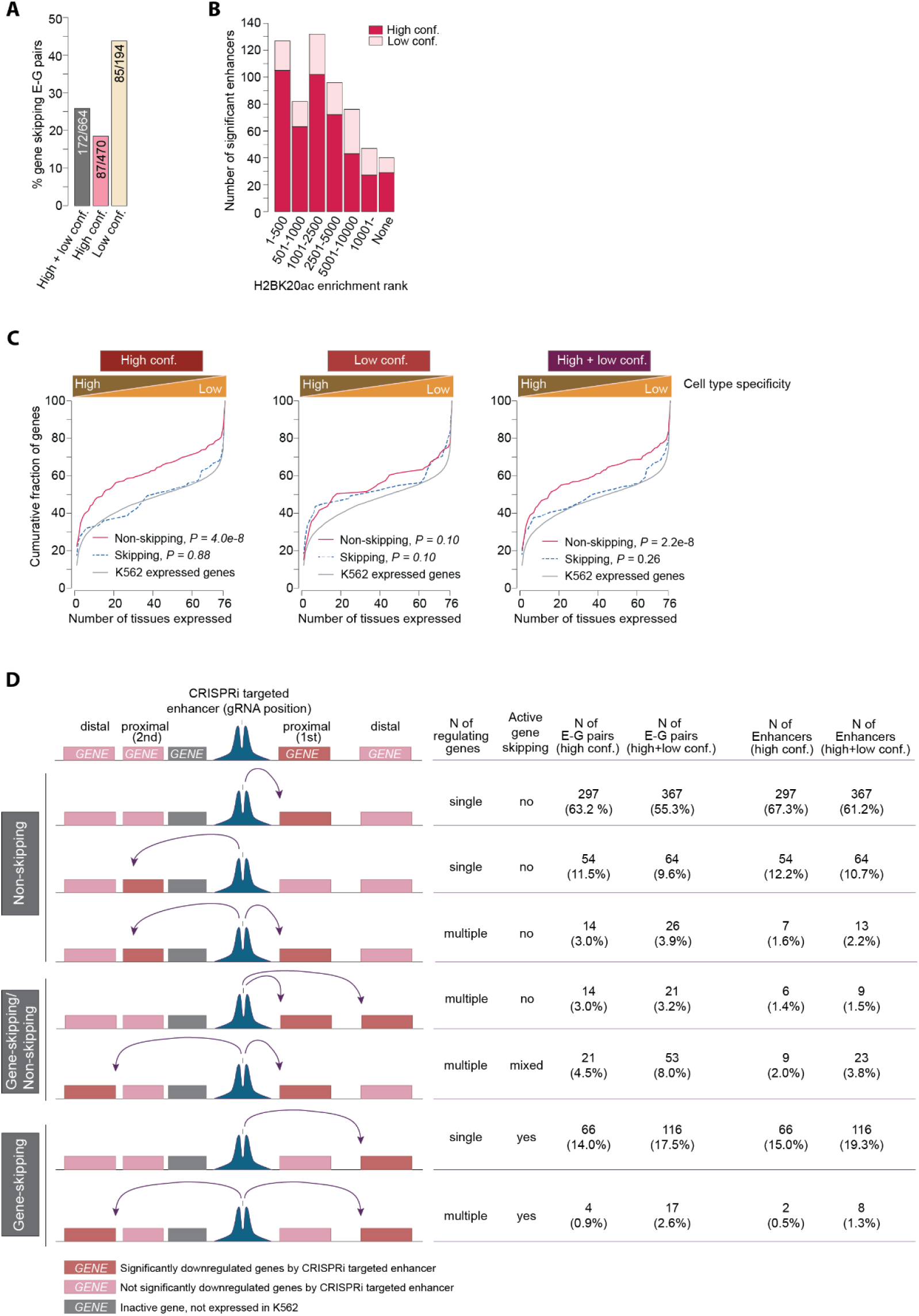
H2BK20ac marks most CRISPRi functional scoring enhancers, and the targets of H2BK20ac-marked enhancers are biased for cell-type-specific genes. (**A**) Fraction of E-G pairs that skip proximal active genes in the specified enhancer confidence groups. (**B**) The number of significant scoring enhancers is shown, categorized into high confidence (High conf.) and low confidence (Low conf.) groups, and sub-grouped in the indicated H2BK20ac enrichment rank class. This analysis includes all 600 functional enhancers identified by Gasperini et al.^40^. Some enhancers are linked to multiple genes and include high-confidence and low-confidence E-G pairs; these are considered high-confidence enhancers in this analysis. (**C**) Non-skipping enhancer targets are biased for cell-type-specific genes. The cumulative fraction of skipping and non-skipping enhancer target genes plotted against genes expressed in 76 different human fetal and adult tissues^46,47^. High-confidence and low-confidence enhancers are classified into skipping and non-skipping categories. Tissue specificity is determined by plotting CRISPRi-identified enhancer target genes against genes expressed in human tissues. The tissue specificity of all genes expressed in K562 is shown as a reference. Except for the low-confidence group, targets of non-skipping enhancers are biased for greater cell-type specificity compared to targets of gene-skipping enhancers. Two-sided Kolmogorov-Smirnov tests, High conf.; gene non-skipping versus expressed genes *P*=4.0X10^-8^, gene skipping versus expressed genes *P*=0.88, low conf.; gene non-skipping versus expressed genes *P*=0.10, gene skipping versus expressed genes *P*=0.10, high conf. + low conf.; gene non-skipping versus expressed genes *P*=2.2X10^-8^, gene skipping versus expressed genes *P*=0.26. (**D**) Within the indicated gene skipping class, the number and fraction of CRISPRi-defined functional enhancers and E-G pairs that involve proximal or distal genes, single or multiple.

**Supplemental Figure 2.**
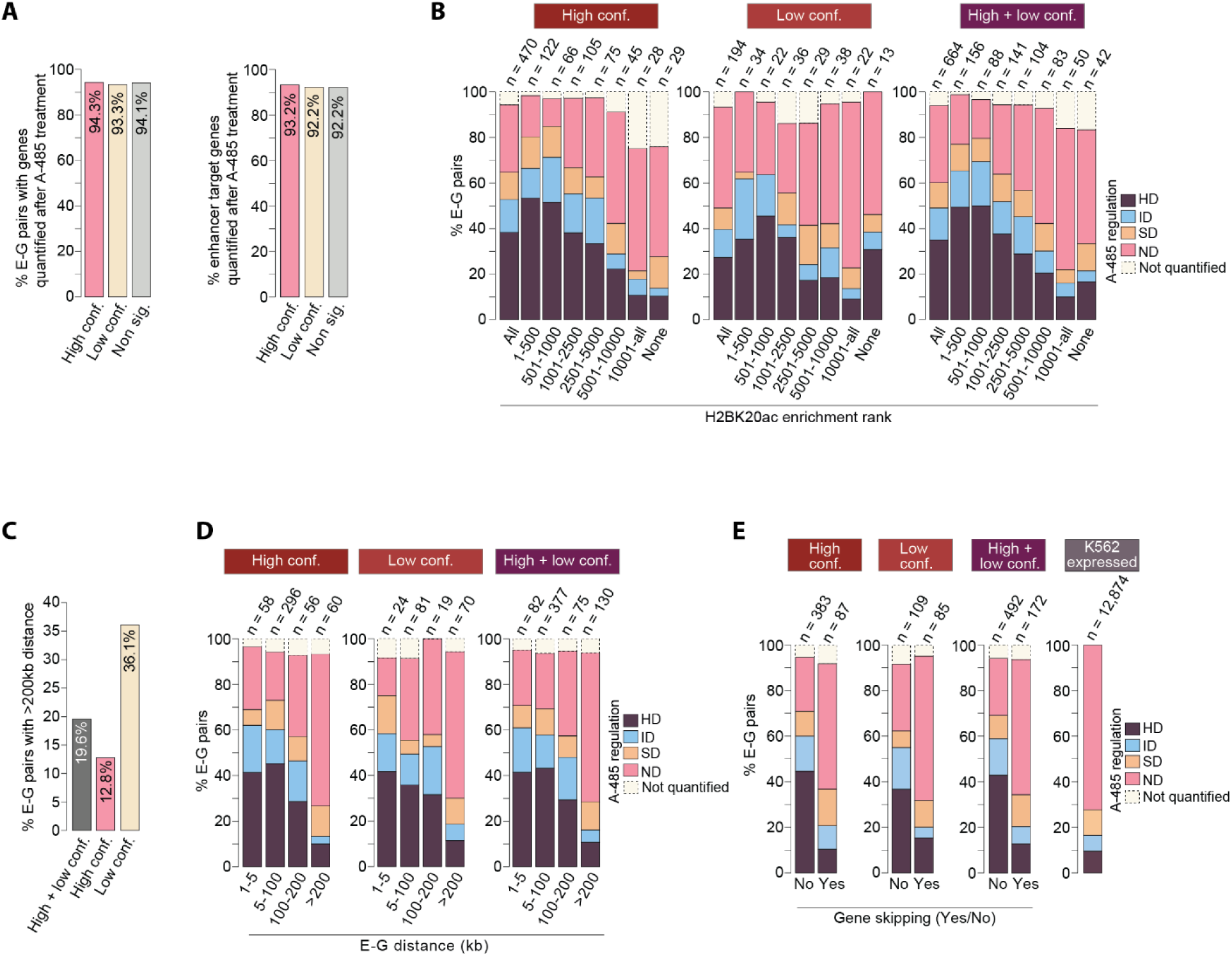
CBP/p300 is preferentially required for activating genes occurring proximally to H2BK20ac marked enhancers. (**A**) The fraction of genes in CRISPRi-defined E-G pairs (left panel) and enhancer target genes (right panel) for which A-485-induced nascent transcription data were available in the Narita et al. dataset^48^. (**B**) In the indicated enhancer confidence groups, E-G pairs are sub-grouped based on enhancer H2BK20ac rank, and the fraction of A-485-regulated genes is shown within the indicated rank categories. NA: gene expression data are not available in A-485 treated cells. (**C**) Fraction of E-G pairs with very long (>200kb) distances in the indicated enhancer confidence groups. (**D**) E-G pairs are classified based on E-G distance in the indicated enhancer confidence groups, and the fraction of A-485-regulated genes is shown within the indicated categories. (**E**) In the indicated enhancer confidence groups, E-G pairs are classified into gene skipping and non-skipping classes, and within each class, the fraction of A-485-regulated genes is depicted.

**Supplemental Figure 3.**
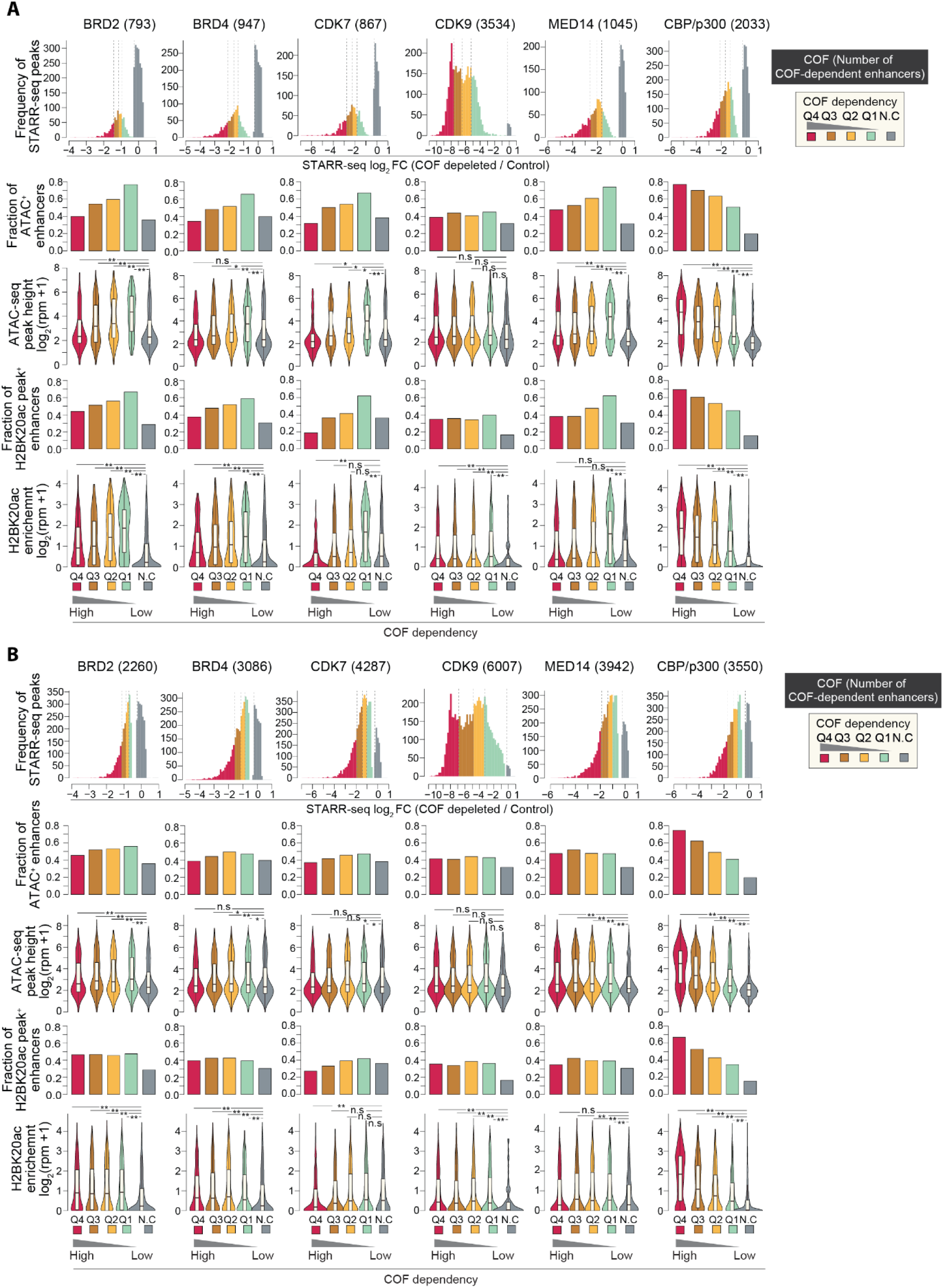
CBP/p300 is required for the function of natively accessible enhancers in STARR-seq. (**A**) The requirement of the six indicated cofactors (COFs; BRD2, BRD4, CDK7, CDK9, MED14, CBP/p300) for enhancer activity was assessed by Neumayr et al. using STARR seq^52^. For each of the analyzed COFs, enhancers showing statistically significant (p=<0.05) COF dependency were grouped into quartiles. The fraction of enhancers showing a decrease in STARR-seq activity after the depletion of the indicated COFs is indicated. Enhancers showing no change (N.C.) in activity after the depletion of the indicated COFs are shown as a separate group. In each of the analyzed COFs, and enhancer groups, H2BK20ac enrichment in enhancers and their ATAC-seq accessibility in native chromatin was determined in the cell line in which the enhancer activity was assayed (HCT116). In the top raw, the frequency plot shows the fold-change distribution of enhancer activity in COF-depleted and wild-type control. In the second row, bar charts show the fraction of enhancers that are ATAC-seq^+^. In the third row, violin plots show DNA accessibility by ATAC-seq. In the fourth raw, bar charts show the fraction of enhancers that are H2BK20ac^+^. In the final row, violin plots show H2BK20ac enrichment. Two-sided Mann–Whitney U-test, adjusted for multiple comparisons with the Benjamini– Hochberg method, ** *P* < 0.001, *P* < 0.05. (**B**) H2BK20ac enrichment and ATAC-seq accessibility of STARR-seq-analyzed enhancers were determined in the same manner as described in panel A, except that in this analysis, include enhancers whose activity in STARR-seq is decreased by >1.5 fold (regardless of statistical significance) in the indicated COF-depleted cells. Note that, among all analyzed COFs, CBP/p300 is the only COF whose requirement for enhancer function is strongly associated with the enrichment of H2BK20ac in enhancers and ATAC-seq accessibility of enhancers in the native

**Supplemental Figure 4 C.**
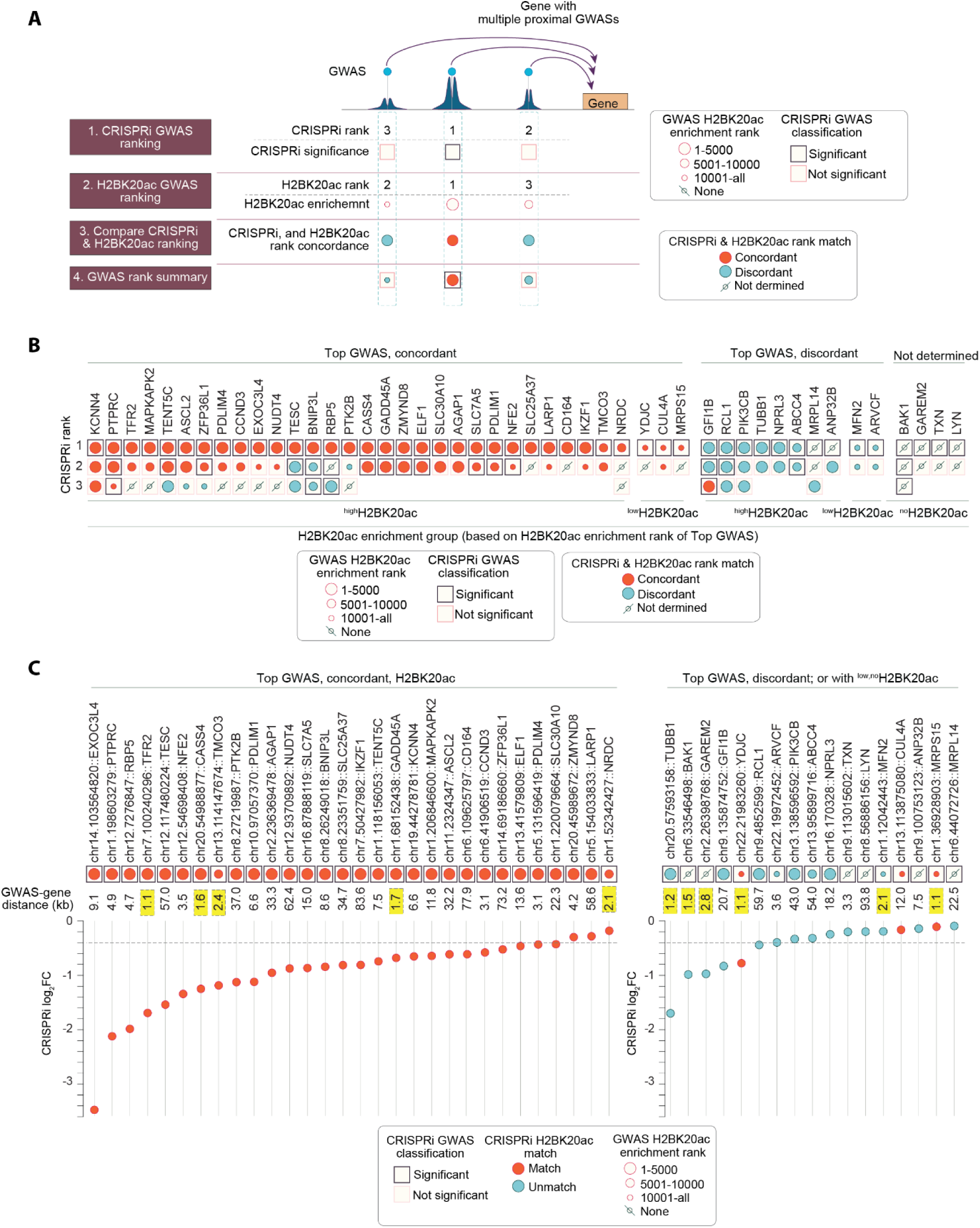
Overlap inRISPRi targeted enhancers and GWAS loci, and H2BK20ac enrichment-based prediction of relative GWAS strength. (**A**) Schematic representation of the strategy for comparing CRISPRi-determined GWAS strength with the H2BK20ac enrichment-predicted GWAS strength. This analysis includes genes (n=47) where CRISPRi targeted multiple proximal GWASs^41^, and CRISPRi targeting of at least one of the targeted GWASs led to significant downregulation of the nearest active gene. The concordance in CRISPRi-measured and H2BK20ac-predicted GWAS functional rank was analyzed as follows: (1) For each gene, proximal GWASs are ranked (strongest 1, intermediate 2, weakest 3) based on CRISPRi-induced downregulation of the GWAS proximal gene. The ranking includes up to 3 GWASs causing strong fold-downregulation by CRISPRi and includes both significant and non-significant scoring GWASs. (2) For each gene, the strength of proximal GWASs is predicted based on the H2BK20ac enrichment. (3) The concordance between CRISPRi-determined and H2BK20ac-predicted GWAS ranks is compared for each gene. (4) GWASs are classified as follows: Concordant GWAS: CRISPRi-determined GWAS rank is the same as the rank predicted by H2BK20ac; Discordant GWAS: CRISPRi-determined GWAS rank is not the same as predicted by H2BK20ac; Not determined: GWAS rank not predicted because none of the proximal GWASs are marked with H2BK20ac. (**B**) The concordance between CRISPRi-defined and H2BK20ac-predicted GWAS ranking for GWASs occurring proximally to the indicated genes is shown. CRISPRi and H2BK20ac-based GWAS ranking was determined as described above in panel A. For each gene, CRISPRi-determined GWAS rank, the significance of gene downregulation in CRISPRi experiments (significant, not significant), H2BK20ac enrichment rank, and concordance in CRISPRi-determined and H2BK20ac-predicted GWAS ranking are depicted. GWASs are sorted from strongest to weakest (#1 to #3) based on CRISPRi-determined GWAS strength. Top GWAS: GWAS causing the strongest downregulation of the proximal gene in CRISPRi. (**C**) Attributes GWASs causing the most robust gene downregulation of the proximal gene in CRISPRi (Top GWAS). The genome coordinates of Top GWASs and their target gene names, GWAS-target gene distance, and CRISPRi-induced fold downregulation of proximal GWAS target genes are shown. The left panel shows GWAS-gene pairs involving ^high^H2BK20ac marked concordant GWASs, while the right panel displays GWAS-gene pairs with ^low,no^H2BK20ac or discordant matching GWASs. Note that discordant matching and ^low,no^H2BK20ac GWASs cause weak gene downregulation in CRISPRi unless occurring close to target genes (<3kb from TSS). This analysis shows that H2BK20ac enrichment can predict the strongest proximal GWAS, except in instances where GWASs have low H2BK20ac enrichment or occur near the promoters of their target genes.

**Supplemental Figure 5.**
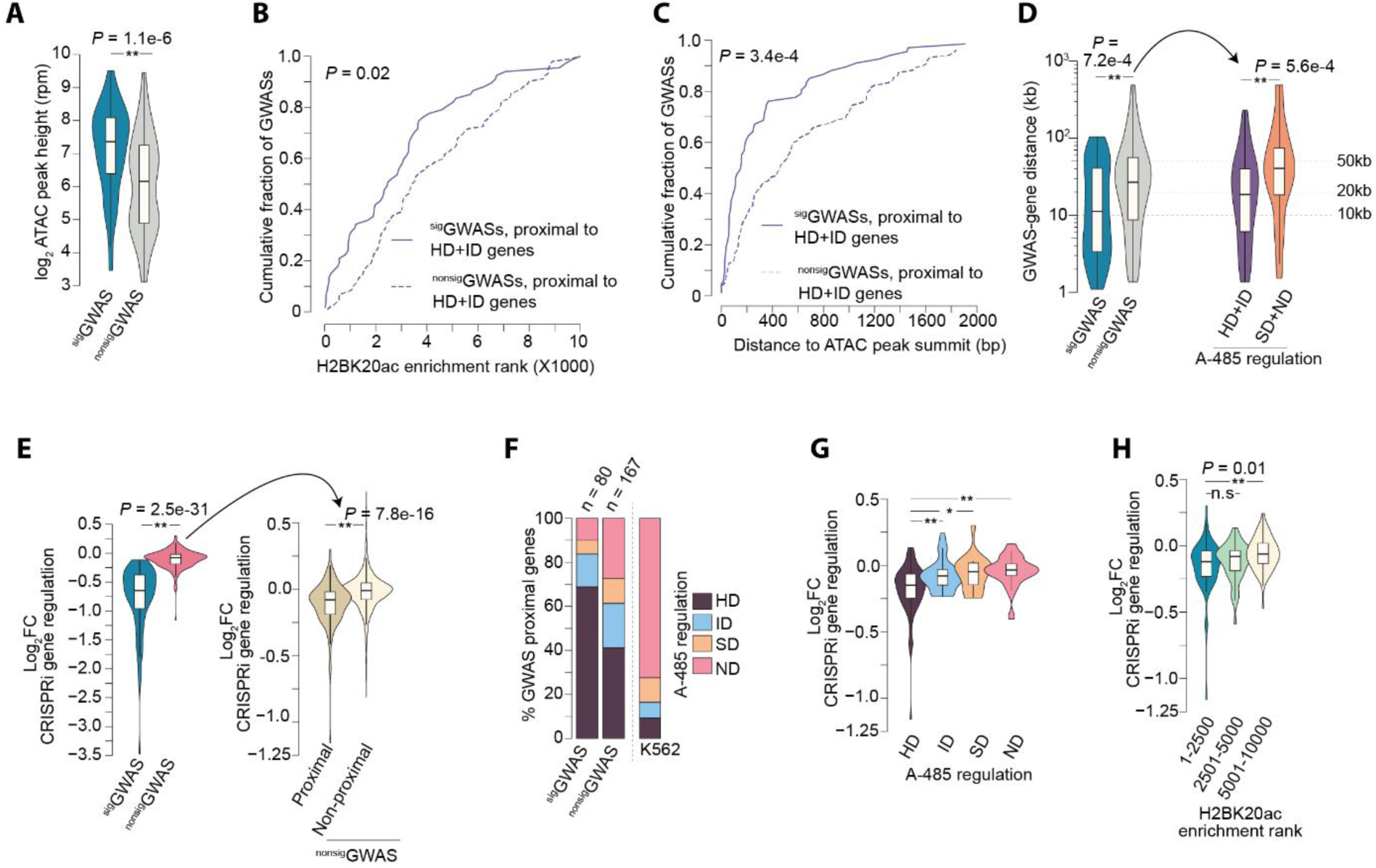
CRISPRi is underpowered to detect weak but potentially functional GWAS loci. (**A**) ATAC peak height significant GWAS (^sig^GWAS) and non-significant GWAS (^nonsig^GWAS) groups. Only ^high^H2BK20ac GWASs within 1kb from the ATAC peak summit were analyzed. ^sig^GWAS are those where CRISPRi targeting led to statistically significant downregulation of their targets, while ^nonsig^GWAS are those that did not cause a significant reduction in target gene expression. This analysis only includes GWASs with H2BK20ac enrichment rank 1-10,000 (^high^H2BK20ac). Two-sided Mann–Whitney U-test, *P*=1.1X10^-6^. (**B-C**) Cumulative plots showing GWAS H2BK20ac enrichment rank (**B**) and distance from ATAC peak summit (**C**) in ^sig^GWASs and ^nonsig^GWASs occurring proximally to A-485 downregulated (HD+ID) genes. Statistical significance was determined by two-sided Kolmogorov-Smirnov tests. (**D**) GWAS-gene distance among ^sig^GWAS and ^nonsig^GWAS groups (left panels). ^nonsig^GWASs are sub-grouped based on the A-485-induced regulation of the GWAS proximal genes (right panel), and GWAS-gene distances for the indicated groups of A-485 downregulated genes are shown. Two-sided Mann–Whitney U-test, *P*=5.6X10^-4^. (**E**) CRISPRi-induced regulation of sigGWAS and nonsigGWAS proximal genes is shown (left panel). Within the ^nonsig^GWAS group, the right panel shows CRISPRi-induced regulation of the most proximal (nearest) and non-proximal genes. Two-sided Mann–Whitney U-test, Sig. versus Non-sig. *P*=2.5X10^-31^, Proximal versus distal *P*=7.8X10^-16^. (**F**) A-485-induced regulation of genes occurring most proximally to ^sig^GWAS and ^nonsig^GWAS groups. A-485-induced regulation of K562-expressed genes is shown as a reference. (**G**) CRISPRi-targeted ^high^H2BK20ac GWASs were grouped based on A-485-induced downregulation of nearest genes, and CRISPRi-induced change in the expression of nearest genes is shown. Two-sided Mann–Whitney U-test, adjusted for multiple comparisons with the Benjamini–Hochberg method; \**P*< 0.05, \*\**P*<0.01. (**H**) CRISPRi-induced downregulation of genes occurring nearest to the indicated GWAS H2BK20ac enrichment ranks groups. Two-sided Mann–Whitney U-test, adjusted for multiple comparisons with the Benjamini–Hochberg method, n.s not significant, \*\**P*<0.01.

**Supplemental Figure 6.**
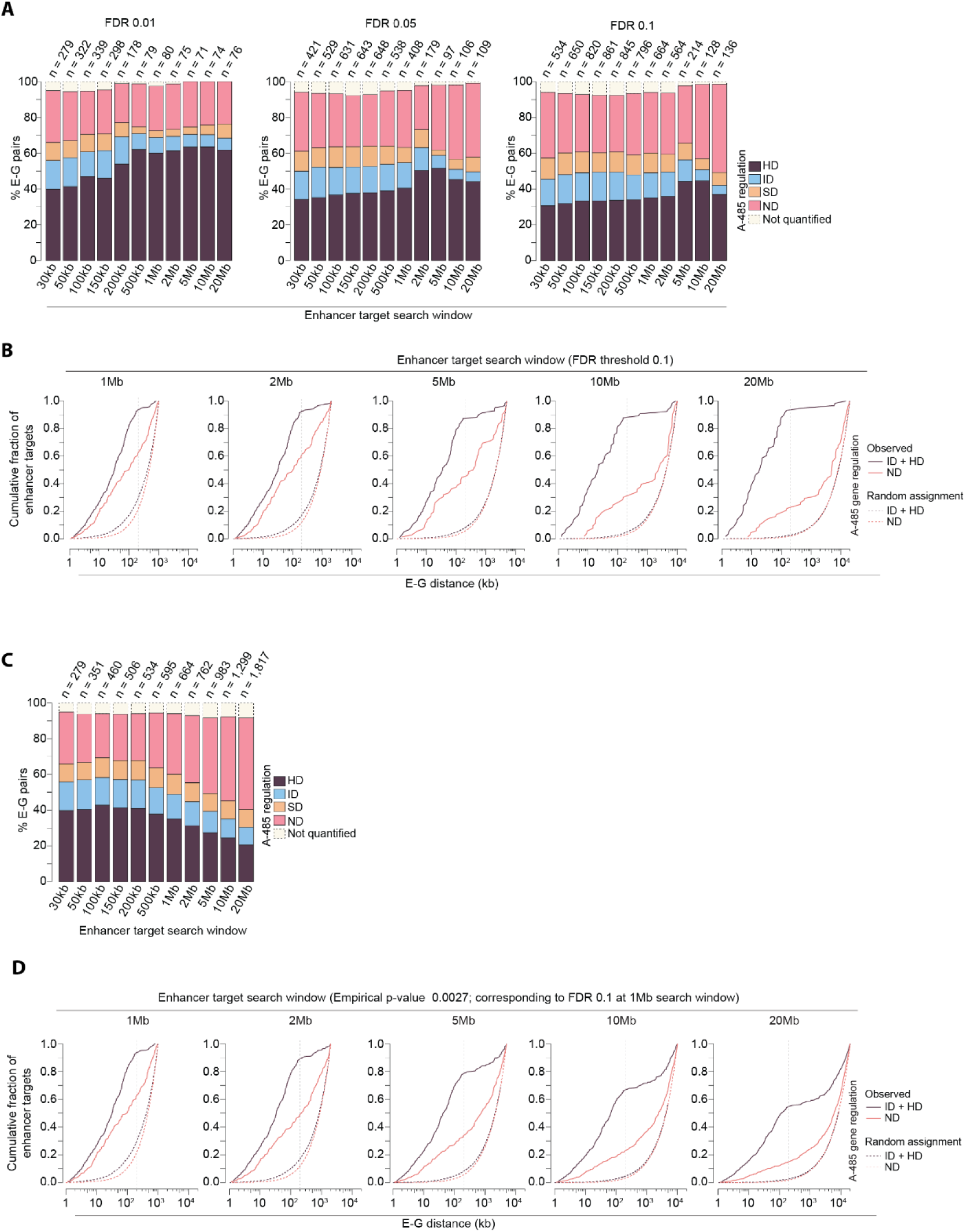
Enhancer target search window in CRISPRi analyses impacts the number of significant scoring enhancers and the nature of enhancer targets identified. (**A**) Shown is the number of significant scoring E-G pairs identified at the indicated search windows and FDRs and the fraction of A-485-regulated genes within each group. Gasperini et al. CRISPRi data^40^ was analyzed using the indicated distance windows at the false discovery rates of 0.1, 0.05, and 0.01. (**B**) The cumulative fraction of E-G pairs involving the indicated groups of A-485-regulated genes in the specified target search windows. E-G pairs were identified at the indicated genomic distance windows at an FDR of 0.1. As a reference, we randomly chose the same number of genes as the number of significant genes within the target search window and categorized the genes based on A-485-induced regulation. Results from multiple iterations (N = 1,000) were combined and used as a reference random group. (**C**) The number of E-G pairs identified at the indicated target search window at a fixed p-value (yielding FDR 0.1 at 1MB target search window), along with the fraction of E-G pairs involving A-485-regulated genes. Of note, broadening the enhancer target search window increases the fraction of enhancers linked to ND genes and decreases the fraction of enhancers linked to HD+ID genes. (**D**) The cumulative fraction of E-G pairs involving the indicated groups of A-485-regulated genes in the specified enhancer target search windows. E-G pairs were identified at the indicated genomic windows at a fixed p-value (yielding FDR 0.1 at 1MB target search window). As a reference, we randomly chose the same number of genes as the number of significant genes within the target search window and categorized the genes based on A-485-induced regulation. Results from multiple iterations (N = 1,000) were combined and used as a reference random group.

## References

1 de Laat, W. & Duboule, D. Topology of mammalian developmental enhancers and their regulatory landscapes. Nature 502, 499–506, doi:10.1038/nature12753 (2013).

2 Spitz, F. & Furlong, E. E. Transcription factors: from enhancer binding to developmental control. Nat Rev Genet 13, 613–626, doi:10.1038/nrg3207 (2012).

3 Shlyueva, D., Stampfel, G. & Stark, A. Transcriptional enhancers: from properties to genome-wide predictions. Nat Rev Genet 15, 272–286, doi:10.1038/nrg3682 (2014).

4 Maurano, M. T. et al. Systematic localization of common disease-associated variation in regulatory DNA. Science 337, 1190–1195, doi:10.1126/science.1222794 (2012).

5 Visscher, P. M. et al. 10 Years of GWAS Discovery: Biology, Function, and Translation. Am J Hum Genet 101, 5–22, doi:10.1016/j.ajhg.2017.06.005 (2017).

6 Uffelmann, E. et al. Genome-wide association studies. Nature Reviews Methods Primers 1, 59, doi:10.1038/s43586-021-00056-9 (2021).

7 Schoenfelder, S. & Fraser, P. Long-range enhancer-promoter contacts in gene expression control. Nat Rev Genet 20, 437–455, doi:10.1038/s41576-019-0128-0 (2019).

8 Dixon, J. R. et al. Topological domains in mammalian genomes identified by analysis of chromatin interactions. Nature 485, 376–380, doi:10.1038/nature11082 (2012).

9 Nora, E. P. et al. Spatial partitioning of the regulatory landscape of the X-inactivation centre. Nature 485, 381–385, doi:10.1038/nature11049 (2012).

10 Rao, S. S. et al. A 3D map of the human genome at kilobase resolution reveals principles of chromatin looping. Cell 159, 1665–1680, doi:10.1016/j.cell.2014.11.021 (2014).

11 Lettice, L. A. et al. A long-range Shh enhancer regulates expression in the developing limb and fin and is associated with preaxial polydactyly. Hum Mol Genet 12, 1725–1735, doi:10.1093/hmg/ddg180 (2003).

12 Bell, A. C. & Felsenfeld, G. Methylation of a CTCF-dependent boundary controls imprinted expression of the Igf2 gene. Nature 405, 482–485, doi:10.1038/35013100 (2000).

13 Oudelaar, A. M., Beagrie, R. A., Kassouf, M. T. & Higgs, D. R. The mouse alpha-globin cluster: a paradigm for studying genome regulation and organization. Curr Opin Genet Dev 67, 18–24, doi:10.1016/j.gde.2020.10.003 (2021).

14 Tolhuis, B., Palstra, R. J., Splinter, E., Grosveld, F. & de Laat, W. Looping and interaction between hypersensitive sites in the active beta-globin locus. Mol Cell 10, 1453–1465, doi:10.1016/s1097-2765(02)00781-5 (2002).

15 Zhang, Y. et al. Chromatin connectivity maps reveal dynamic promoter-enhancer long-range associations. Nature 504, 306–310, doi:10.1038/nature12716 (2013).

16 Hsieh, T. S. et al. Resolving the 3D Landscape of Transcription-Linked Mammalian Chromatin Folding. Mol Cell 78, 539–553 e538, doi:10.1016/j.molcel.2020.03.002 (2020).

17 Li, G. et al. Extensive promoter-centered chromatin interactions provide a topological basis for transcription regulation. Cell 148, 84–98, doi:10.1016/j.cell.2011.12.014 (2012).

18 Sanyal, A., Lajoie, B. R., Jain, G. & Dekker, J. The long-range interaction landscape of gene promoters. Nature 489, 109–113, doi:10.1038/nature11279 (2012).

19 Chandra, V. et al. Promoter-interacting expression quantitative trait loci are enriched for functional genetic variants. Nat Genet 53, 110–119, doi:10.1038/s41588-020-00745-3 (2021).

20 Winick-Ng, W. et al. Cell-type specialization is encoded by specific chromatin topologies. Nature 599, 684–691, doi:10.1038/s41586-021-04081-2 (2021).

21 Tomas-Daza, L. et al. Low input capture Hi-C (liCHi-C) identifies promoter-enhancer interactions at high-resolution. Nat Commun 14, 268, doi:10.1038/s41467-023-35911-8 (2023).

22 Beagrie, R. A. et al. Complex multi-enhancer contacts captured by genome architecture mapping. Nature 543, 519–524, doi:10.1038/nature21411 (2017).

23 Mumbach, M. R. et al. Enhancer connectome in primary human cells identifies target genes of disease-associated DNA elements. Nat Genet 49, 1602–1612, doi:10.1038/ng.3963 (2017).

24 Javierre, B. M. et al. Lineage-Specific Genome Architecture Links Enhancers and Non-coding Disease Variants to Target Gene Promoters. Cell 167, 1369–1384 e1319, doi:10.1016/j.cell.2016.09.037 (2016).

25 Lu, L. et al. Robust Hi-C Maps of Enhancer-Promoter Interactions Reveal the Function of Non-coding Genome in Neural Development and Diseases. Mol Cell 79, 521–534 e515, doi:10.1016/j.molcel.2020.06.007 (2020).

26 Dowen, J. M. et al. Control of cell identity genes occurs in insulated neighborhoods in mammalian chromosomes. Cell 159, 374–387, doi:10.1016/j.cell.2014.09.030 (2014).

27 Vangala, P. et al. High-Resolution Mapping of Multiway Enhancer-Promoter Interactions Regulating Pathogen Detection. Mol Cell 80, 359–373 e358, doi:10.1016/j.molcel.2020.09.005 (2020).

28 Liang, L. et al. Complementary Alu sequences mediate enhancer-promoter selectivity. Nature, doi:10.1038/s41586-023-06323-x (2023).

29 Tan, L. et al. Changes in genome architecture and transcriptional dynamics progress independently of sensory experience during post-natal brain development. Cell 184, 741–758 e717, doi:10.1016/j.cell.2020.12.032 (2021).

30 Lando, D. et al. Enhancer-promoter interactions are reconfigured through the formation of long-range multiway hubs as mouse ES cells exit pluripotency. Mol Cell, doi:10.1016/j.molcel.2024.02.015 (2024).

31 Liu, Z. et al. Linking genome structures to functions by simultaneous single-cell Hi-C and RNA-seq. Science 380, 1070–1076, doi:10.1126/science.adg3797 (2023).

32 Monahan, K., Horta, A. & Lomvardas, S. LHX2- and LDB1-mediated trans interactions regulate olfactory receptor choice. Nature 565, 448–453, doi:10.1038/s41586-018-0845-0 (2019).

33 Oudelaar, A. M. et al. Single-allele chromatin interactions identify regulatory hubs in dynamic compartmentalized domains. Nat Genet 50, 1744–1751, doi:10.1038/s41588-018-0253-2 (2018).

34 Goel, V. Y., Huseyin, M. K. & Hansen, A. S. Region Capture Micro-C reveals coalescence of enhancers and promoters into nested microcompartments. Nat Genet 55, 1048–1056, doi:10.1038/s41588-023-01391-1 (2023).

35 Aboreden, N. G. et al. LDB1 establishes multi-enhancer networks to regulate gene expression. bioRxiv, 2024.2008.2023.609430, doi:10.1101/2024.08.23.609430 (2024).

36 Beagrie, R. A. et al. Multiplex-GAM: genome-wide identification of chromatin contacts yields insights overlooked by Hi-C. Nat Methods 20, 1037–1047, doi:10.1038/s41592-023-01903-1 (2023).

37 Wen, X. et al. Single-cell multiplex chromatin and RNA interactions in ageing human brain. Nature 628, 648–656, doi:10.1038/s41586-024-07239-w (2024).

38 Wu, H. et al. Simultaneous single-cell three-dimensional genome and gene expression profiling uncovers dynamic enhancer connectivity underlying olfactory receptor choice. Nat Methods 21, 974–982, doi:10.1038/s41592-024-02239-0 (2024).

39 Zhou, T. et al. GAGE-seq concurrently profiles multiscale 3D genome organization and gene expression in single cells. Nat Genet, doi:10.1038/s41588-024-01745-3 (2024).

40 Gasperini, M. et al. A Genome-wide Framework for Mapping Gene Regulation via Cellular Genetic Screens. Cell 176, 377–390 e319, doi:10.1016/j.cell.2018.11.029 (2019).

41 Morris, J. A. et al. Discovery of target genes and pathways at GWAS loci by pooled single-cell CRISPR screens. Science 380, eadh7699, doi:10.1126/science.adh7699 (2023).

42 Barshad, G. et al. RNA polymerase II dynamics shape enhancer-promoter interactions. Nat Genet 55, 1370–1380, doi:10.1038/s41588-023-01442-7 (2023).

43 Whyte, W. A. et al. Master transcription factors and mediator establish super-enhancers at key cell identity genes. Cell 153, 307–319, doi:10.1016/j.cell.2013.03.035 (2013).

44 Krietenstein, N. et al. Ultrastructural Details of Mammalian Chromosome Architecture. Mol Cell 78, 554–565 e557, doi:10.1016/j.molcel.2020.03.003 (2020).

45 Durand, N. C. et al. Juicer Provides a One-Click System for Analyzing Loop-Resolution Hi-C Experiments. Cell Syst 3, 95–98, doi:10.1016/j.cels.2016.07.002 (2016).

46 Yu, N. Y. et al. Complementing tissue characterization by integrating transcriptome profiling from the Human Protein Atlas and from the FANTOM5 consortium. Nucleic Acids Res 43, 6787–6798, doi:10.1093/nar/gkv608 (2015).

47 Hon, C. C. et al. An atlas of human long non-coding RNAs with accurate 5’ ends. Nature 543, 199–204, doi:10.1038/nature21374 (2017).

48 Narita, T. et al. Acetylation of histone H2B marks active enhancers and predicts CBP/p300 target genes. Nat Genet 55, 679–692, doi:10.1038/s41588-023-01348-4 (2023).

49 Weinert, B. T. et al. Time-Resolved Analysis Reveals Rapid Dynamics and Broad Scope of the CBP/p300 Acetylome. Cell 174, 231–244 e212, doi:10.1016/j.cell.2018.04.033 (2018).

50 Narita, T. et al. The logic of native enhancer-promoter compatibility and cell-type-specific gene expression variation. bioRxiv, 2022.2007.2018.500456, doi:10.1101/2022.07.18.500456 (2022).

51 Lasko, L. M. et al. Discovery of a selective catalytic p300/CBP inhibitor that targets lineage-specific tumours. Nature 550, 128–132, doi:10.1038/nature24028 (2017).

52 Neumayr, C. et al. Differential cofactor dependencies define distinct types of human enhancers. Nature 606, 406–413, doi:10.1038/s41586-022-04779-x (2022).

53 Peng, T. et al. STARR-seq identifies active, chromatin-masked, and dormant enhancers in pluripotent mouse embryonic stem cells. Genome Biol 21, 243, doi:10.1186/s13059-020-02156-3 (2020).

54 Sahu, B. et al. Sequence determinants of human gene regulatory elements. Nat Genet 54, 283–294, doi:10.1038/s41588-021-01009-4 (2022).

55 Mannion, B. J. et al. Uncovering Hidden Enhancers Through Unbiased <em>In Vivo</em> Testing. bioRxiv, 2022.2005.2029.493901, doi:10.1101/2022.05.29.493901 (2022).

56 Akgol Oksuz, B., et al. Systematic evaluation of chromosome conformation capture assays. Nat Methods 18, 1046–1055, doi:10.1038/s41592-021-01248-7 (2021).

57 Moorthy, S. D. et al. Enhancers and super-enhancers have an equivalent regulatory role in embryonic stem cells through regulation of single or multiple genes. Genome Res 27, 246–258, doi:10.1101/gr.210930.116 (2017).

58 Pradeepa, M. M. et al. Histone H3 globular domain acetylation identifies a new class of enhancers. Nat Genet 48, 681–686, doi:10.1038/ng.3550 (2016).

59 Kumar, V. et al. Comprehensive benchmarking reveals H2BK20 acetylation as a distinctive signature of cell-state-specific enhancers and promoters. Genome Res 26, 612–623, doi:10.1101/gr.201038.115 (2016).

60 Regadas, I. et al. A unique histone 3 lysine 14 chromatin signature underlies tissue-specific gene regulation. Mol Cell 81, 1766–1780 e1710, doi:10.1016/j.molcel.2021.01.041 (2021).

61 Hua, P. et al. Defining genome architecture at base-pair resolution. Nature 595, 125–129, doi:10.1038/s41586-021-03639-4 (2021).

62 Haberle, V. et al. Transcriptional cofactors display specificity for distinct types of core promoters. Nature 570, 122–126, doi:10.1038/s41586-019-1210-7 (2019).

63 Narita, T., et al. A unified model of gene expression control by cohesin and CTCF. BioRxiv (2024).

64 Hilton, I. B. et al. Epigenome editing by a CRISPR-Cas9-based acetyltransferase activates genes from promoters and enhancers. Nat Biotechnol 33, 510–517, doi:10.1038/nbt.3199 (2015).

65 Zuin, J. et al. Nonlinear control of transcription through enhancer-promoter interactions. Nature 604, 571–577, doi:10.1038/s41586-022-04570-y (2022).

66 Zhernakova, D. V. et al. Identification of context-dependent expression quantitative trait loci in whole blood. Nat Genet 49, 139–145, doi:10.1038/ng.3737 (2017).

67 Strober, B. J. et al. Dynamic genetic regulation of gene expression during cellular differentiation. Science 364, 1287–1290, doi:10.1126/science.aaw0040 (2019).

68 Kim-Hellmuth, S. et al. Cell type-specific genetic regulation of gene expression across human tissues. Science 369, doi:10.1126/science.aaz8528 (2020).

69 Mostafavi, H., Spence, J. P., Naqvi, S. & Pritchard, J. K. Limited overlap of eQTLs and GWAS hits due to systematic differences in discovery. bioRxiv, 2022.2005.2007.491045, doi:10.1101/2022.05.07.491045 (2022).

70 Vosa, U. et al. Large-scale cis- and trans-eQTL analyses identify thousands of genetic loci and polygenic scores that regulate blood gene expression. Nat Genet 53, 1300–1310, doi:10.1038/s41588-021-00913-z (2021).

71 Kreibich, E. & Krebs, A. R. Cofactors: a new layer of specificity to enhancer regulation. Trends Biochem Sci 47, 993–995, doi:10.1016/j.tibs.2022.07.008 (2022).

72 Li, H. & Durbin, R. Fast and accurate short read alignment with Burrows-Wheeler transform. Bioinformatics 25, 1754–1760, doi:10.1093/bioinformatics/btp324 (2009).

73 Li, H. et al. The Sequence Alignment/Map format and SAMtools. Bioinformatics 25, 2078–2079, doi:10.1093/bioinformatics/btp352 (2009).

74 Quinlan, A. R. & Hall, I. M. BEDTools: a flexible suite of utilities for comparing genomic features. Bioinformatics 26, 841–842, doi:10.1093/bioinformatics/btq033 (2010).

75 Anders, S., Pyl, P. T. & Huber, W. HTSeq--a Python framework to work with high-throughput sequencing data. Bioinformatics 31, 166–169, doi:10.1093/bioinformatics/btu638 (2015).

76 Love, M. I., Huber, W. & Anders, S. Moderated estimation of fold change and dispersion for RNA-seq data with DESeq2. Genome Biol 15, 550, doi:10.1186/s13059-014-0550-8 (2014).

77 Dobin, A. et al. STAR: ultrafast universal RNA-seq aligner. Bioinformatics 29, 15–21, doi:10.1093/bioinformatics/bts635 (2013).

78 Hentges, L. D. et al. LanceOtron: a deep learning peak caller for genome sequencing experiments. Bioinformatics 38, 4255–4263, doi:10.1093/bioinformatics/btac525 (2022).

79 Ramirez, F. et al. deepTools2: a next generation web server for deep-sequencing data analysis. Nucleic Acids Res 44, W160–165, doi:10.1093/nar/gkw257 (2016).

80 Jung, Y. & Han, D. BWA-MEME: BWA-MEM emulated with a machine learning approach. Bioinformatics 38, 2404–2413, doi:10.1093/bioinformatics/btac137 (2022).

81 Khan, A. & Zhang, X. dbSUPER: a database of super-enhancers in mouse and human genome. Nucleic Acids Res 44, D164–171, doi:10.1093/nar/gkv1002 (2016).

82 Kent, W. J. et al. The human genome browser at UCSC. Genome Res 12, 996–1006, doi:10.1101/gr.229102 (2002).

83 Narita, T. et al. Enhancers are activated by p300/CBP activity-dependent PIC assembly, RNAPII recruitment, and pause release. Mol Cell 81, 2166–2182 e2166, doi:10.1016/j.molcel.2021.03.008 (2021).

